# Incorporating Surfaced-Induced Dissociation Mass Spectrometry Data into an AlphaFold-derived deep learning network improves protein structure prediction

**DOI:** 10.64898/2026.06.26.734850

**Authors:** Robert M. Bolz, Elijah H. Day, Zachary C. Drake, Sophie R. Harvey, Vicki H. Wysocki, Steffen Lindert

## Abstract

Surface-Induced Dissociation native Mass Spectrometry (SID-nMS) is a tandem MS activation method that yields information on the connectivity and stoichiometry of protein complexes. While insufficient for direct structure elucidation, the data derived from SID-nMS has considerable potential to inform multimeric protein structure prediction. We hypothesized that incorporating this data into a machine-learning framework could improve multimer prediction accuracy beyond that of existing deep-learning methods. To this end, we developed SIDFold, a novel AlphaFold-based deep-learning network. SIDFold is the first AlphaFold-like network to leverage experimental data during protein complex prediction, and the first deep-learning network to utilize nMS data for structure prediction. We benchmarked SIDFold on the BETA protein set, and observed an improvement in RMSD in 138 of 227 cases including 27 targets in which the predicted structure attained near-native accuracy. We then evaluated the network on 20 proteins with experimental SID-nMS data, yielding an improved RMSD in 18 cases, with five of these cases improving to a high-accuracy complex. Finally, we tested SIDFold against a previously published SID-guided Rosetta docking method, where we saw improvement in 13 of 16 proteins. SIDFold is freely available on GitHub, with example files and commands available in the Supplementary Information.

## Introduction

Protein-protein interactions (PPIs) are crucial components of biological processes. Dysregulation of PPI’s cause several diseases including cancer^1^, arrhythmogenesis^2^, Alzheimer’s^3^, and Parkinson’s^4^, among others. Alongside its structure, the interactome of a protein is a critical determinant of its final function ^5^. Thus, elucidating the structure of interacting protein complexes is crucial for understanding disease mechanisms and developing potential therapeutic interventions^6^. Methods like X-ray Crystallography (XRC)^6^, Nuclear Magnetic Resonance (NMR)^7^, and Cryogenic Electron Microscopy (Cryo-EM)^8^, serve as the ‘gold standard’ for protein structure determination. When successfully applied, all can resolve structure at the atomic level in certain protein systems^9^. For each technique, however, there can be limitations including sample preparation or system requirements restricting their applicability for use in many cases^10,11^.

For the protein systems inaccessible to ‘gold standard’ methods, native Mass Spectrometry (nMS) represents an intriguing alternative to determining structural information^12^. nMS does not provide atomic resolution like the methods described above ^12^. However, it is faster, has low sample requirements (with respect to material used), can handle heterogeneous samples, and has a diverse array of collection methods^12^. In aggregate, this makes nMS a powerful tool for understanding countless protein systems. Different activation methods offer additional, unique insights into protein structure and dynamics. Surface-Induced Dissociation (SID) nMS is an activation method that accelerates a protein complex into a collision surface^13^. This collision transfers the kinetic energy from acceleration into internal, vibrational energy in the protein complex^13^. This internal energy can cause noncovalent dissociation of the protein complex into intact, fragment subcomplexes. This provides information on the stoichiometry, interface size and interface strength for a given protein complex.

The weakest interfaces of a protein complex will break before the stronger interfaces. If the protein is collided at a range of acceleration energies, a dissociation pattern characteristic to the protein complex can be obtained^13^. The short time scale of the technique leads to substructure fragments, suggesting that SID data offers insight into the native structure of the complex^13^. The characteristic dissociation curve shows the relative abundance of each sub-complex for a whole multimeric protein at each acceleration energy. This data is commonly referred to as Energy Resolved Mass Spectrometry (ERMS) data.

SID-nMS data is low-resolution (i.e., yielding no atomic structural details), making it insufficient to provide a full protein structure alone. To obtain a protein structure from this data, an integrative modeling approach must be used. Combining sparse experimental mass spectrometry data with computational methods has a long tradition of successfully elucidating experimentally-derived structures^14–38^. Notably, preliminary work has shown that is possible to model protein structure from the Appearance Energy (AE) in SID-nMS data^39–41^. We subsequently developed a method for simulating ERMS data from a protein structure, given that ERMS data is more information rich than the single point AE data^42^. Most recently, we developed a modeling application in Rosetta^43^ for integrating SID-nMS data into structure prediction^44^. This method used the SID-nMS data to determine the structure of the whole complex given a known monomer structure. This was more accurate than default Rosetta complex modeling. However, while there was a clear improvement over unguided docking, there existed inherent limitations to the method. First, this method was applicable only in cases where the monomeric structure of the protein was known, but not the multimeric structure. And secondly, this method rescored existing multimeric structures, instead of generating them *de novo*. This is a serious limitation in cases where Rosetta does not generate near-native structures for rescoring. Here, we sought to overcome these limitations by creating a new method that generated *de novo* structures without knowing the monomeric structure prior to prediction. This method is the first to directly implement experimental multimer interface data in the *de novo* generation of complex structures.

The Nobel prize winning method, AlphaFold^45^ established a new standard in computational modeling. Since its release, numerous machine learning-based methods have come in its wake^22,46,47^. AlphaFold is widely considered the gold standard in computational modeling. Benchmarks show AlphaFold is on average 76.7% accurate for monomeric structures^48^. However, AlphaFold’s multimeric predictions capabilities lag significantly behind this metric. Independent analysis showed that, on average, AlphaFold’s multimeric predictions are 50% accurate (measured through DockQ_i_), with the accuracy decreasing with increasing oligomeric states^49^. 86% of proteins form a complex in vivo^50^, meaning there is a significant need for accurate multimeric structure prediction.

Networks like AlphaLink^22^ demonstrated the potential for integrating sparse experimental data into deep learning programs. AlphaLink was based on OpenFold, an open source recreation of AlphaFold2 in Pytorch^46^. We hypothesized that integration of SID-nMS data could overcome the limitations of AlphaFold and that an integrated deep learning method would also overcome the above-mentioned limitations of Rosetta-based algorithms, hence improving multimeric structure prediction beyond current capabilities.

In this work, we present SIDFold, the first deep learning network to integrate SID-nMS data into protein structure prediction. This is the first deep learning network of its kind to utilize non-residue level multimeric interface data, which is directly guiding the predictions of conformations and interfaces of the protein complex using experimental data. The network was trained on a curated set of DIPS-Plus proteins specifically chosen for training protein interface predictions^51^. We subset this data to train on structures that are likely candidates for SID-nMS data collection, to better drive the network towards experimentally verified structures. To simulate the SID-ERMS training data, we used a PyRosetta^52^ application^53^. We integrated this application directly into the network. A novel auxiliary loss function fine-tuned the network based on starting AlphaFold2 weights to predict structures that align with the experimental SID data. We benchmarked this network on two protein sets. The first was a benchmark set of 227 proteins from the DIPS-Plus set using simulated SID-ERMS data. We saw improved accuracy in 138/227 proteins, as defined by a decreased Root Mean Square Deviation (RMSD). We then evaluated the network using 20 proteins with experimental SID-ERMS data. In inference, we saw improved multimeric structure accuracy in 18/20 cases, with 15/20 multimeric structures being high accuracy structures (below 5 Å RMSD). Additionally, we also observed improved performance in RMSD_100_, TM-score, DockQ_i_, and interface SASA. This highlights the ability of SIDFold to outperform standard AlphaFold, as well as traditional SID-guided methods to achieve improved multimeric structure prediction.

## Methods

### Generation of training and validation dataset

We trained the network on the DIPS-Plus dataset, a curated set of multimeric structures specifically made for machine learning network training^51^. The ‘raw’ dataset was processed and pruned using the default parameters described in the original publication^51^. Structures were retained only if they had an XRC or Cryo-EM resolution of 3.5 Å or better, contained no protein chain sharing greater than 30% sequence identity to any protein in the Docking Benchmark 5 (DB5) dataset, were comprised of chains of at least 50 residues, and exhibited at least 500 Å^2^ of buried surface area across the whole complex. After filtering, this left the 33,159 protein complexes originally contained in the DIPS-Plus set. Additional filtering criteria were applied to generate the training, validation, and benchmark datasets. Complexes were required to contain between two and eight chains, with no single chain containing more than 150 residues. To limit large differences between subunit size, the number of residues in any two chains in a complex could not vary by more than 10%. Complexes were also required to contain at least 60% non-coil secondary structure according to the Define Secondary Structure of Proteins (DSSP) algorithm^54^, and have a valid symmetry point group annotation deposited in the Protein Data Bank (PDB)^55^. These structural filters selected proteins that are likely candidates for SID-nMS data determination. These proteins are well ordered, have defined symmetry groups, and do not have large size discrepancies between subunits.

The final count of multimeric proteins after this strict filtering requirement was 234. Stringent filtering and training on a smaller set of proteins was inspired by previous works which showed that highly curated experimental structure sets can train a network that outperforms the default DIPS-Plus in accuracy of the network output^56–58^. We randomly split this set into a training set of 200 proteins and a validation set of 34 proteins (to create a randomly generated 85/15 split). The protein IDs and oligomeric state of each protein in the training and validation sets are included in csv format in the supplementary data.

### Generation of benchmark set with BETA proteins and experimental evaluation set

To create a benchmark set, we used the BETA dataset comprised of proteins deposited into the PDB after the AlphaFold templates were selected^59^. This was to ensure the benchmark set was orthogonal to the training and validation sets, and also not include any proteins that AlphaFold was trained on. We used the iteracting_chains_202103_template dataset, and applied the same structural filters that were applied to our training and validation sets (at least 60% ordered, between 50-150 residues per chain, chain lengths within 90%, and between dimers to hexamers). This left 227 proteins, which we used as our benchmark set. Finally, we used the 20 proteins for which we have experimental SID-nMS data as an experimental benchmark set^44^. The protein IDs and oligomeric state of each protein in the benchmark and experimental validation sets are included in csv format in the supplementary data.

### Generation of Multiple Sequence Alignments (MSAs)

Multiple sequence alignments (MSAs) were generated using the default AlphaFold2 workflow ^45^. Briefly, sequence searches and alignment generation were performed using JackHMMER^60^, HHblits^61^, HMMER^62^, and Kalign^63^. The databases searched included the AlphaFold BFD database^45^, the UniRef30 and UniRef90^64^ clustered databases (release 2021_03), MGnify (release 2022_05)^65^, the UniProt^66^ sequence database, and the PDB seqres and PDB 70^55^ clustered sequence databases. For protein complexes, alignments were generated independently for each chain and concatenated during training and inference.

### Connectivity file generation for Training, validation, benchmark, and experimental benchmark datasets

During an SID experiment, a multimeric protein’s symmetry point group is a determining factor in its breakage pattern along with interface properties and can routinely be deduced from the MS data. For this reason, knowledge of the complex’s point group is necessary information to simulate ERMS data from its structure. The symmetry point group of each complex was determined using the QuatSymm implementation provided in the Protein Data Bank GitHub repository ^67,68^. Point group assignments for each protein complex in the training, validation, benchmark, and experimental benchmark datasets are included in the Supplementary Information.

The assigned symmetry point groups were subsequently used to generate the connectivity files required for ERMS data simulation. Each connectivity file specifies the interfaces present in a multimeric complex and identifies interfaces that are equivalent based on the symmetry point group of the complex. Symmetry-equivalent interfaces are assigned an equal probability of dissociation during data simulation. For example, a C4 tetramer contains four symmetric interfaces, A_B, B_C, C_D, and D_A, which are all assigned equal dissociation probabilities. For complexes with no internal symmetry, all interfaces are assumed to be nonequivalent and the strength of each interface is calculated individually during ERMS simulation. For CN complexes (where N equals the number of chains, and N > 1), all interfaces were assumed to be equal during data simulation. For CN complexes where N is less than the number of chains (for example, a C2 tetramer like PDB ID 1WBJ in our experimental benchmark set), the interface areas were calculated and interfaces with areas within 90% of each other were assumed to be equal. For DN complexes, where N is less than the number of chains (for example, a D2 tetramer like PDB ID 1SWB in our experimental benchmark set), the interface areas were also calculated and interfaces with areas within 90% of each other were assumed to be equal. Example connectivity files for several symmetry point groups and code for file generation can be found in SI Table S1 and Supplementary Information.

### Simulation of SID-ERMS data for training, evaluation, and benchmark sets

Experimental SID-ERMS data were not available for the proteins included in the training, evaluation, and benchmark datasets. Therefore, we simulated the SID-ERMS data using the previously published PyRosetta `SID_ERMS_Prediction` application^53^. Data simulation has been used extensively in other deep learning networks during training and validation, producing near-native structures in accordance with experimental data^22,69–71^. The SID-ERMS data was calculated using the connectivity files created above to determine the symmetry to simulate.

### Incorporation of an auxiliary loss function into OpenFold network

The SID-ERMS auxiliary loss function was incorporated into the main recycling loop of the structure module. During training, the 3D coordinates generated during the final recycling iteration were written to the output directory as an intermediate PDB structure. ERMS simulation was performed only for the final recycling iteration because the auxiliary loss function is only evaluated after the completion of the main recycling loop. Simulating ERMS data during earlier iterations would therefore not contribute to the calculated loss and would substantially increase the computational cost of training. Before ERMS simulation, the intermediate structure was refined using a PyRosetta implementation of the Rosetta `FastRelax`^72^ protocol, as side-chain refinement was previously shown to significantly improve the accuracy of simulated ERMS data^42^. To reduce the computational cost of this step, the maximum number of refinement cycles was set to 100. SID-ERMS data was simulated using the relaxed intermediate structure and compared with the corresponding ground truth data by calculating the root-mean-square error (RMSE). For cases where the predicted intermediate structure is inaccurate to the point where there is no interface between the monomer chains, no ERMS simulation is possible and the RMSE was set to 1.0 (Equation 1). This number was chosen because the ERMS is given in fraction of multimeric states, therefore the data ranges from 0.0-1.0. Therefore, an RMSE of 1.0 is not mathematically possible for actual simulated data. The penalty of 1.0 gave the network differentiability between protein complexes without interfaces between them, and protein complexes that have an inaccurate interface. The RMSE is used as the input to the auxiliary loss function, which multiplies the RMSE by the ERMS loss weight, set to 0.01 during training, and incorporated into the total training loss as defined in Equations 2 and 3.

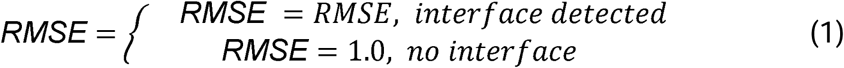

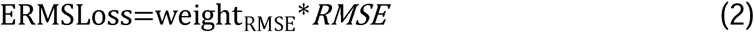

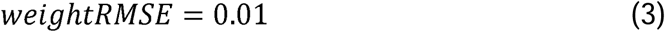

### Training parameters

The training parameters were configured using the DeepSpeed optimization library in PyTorch Lightning^73^. The learning rate was set to 0.001, with ZeRO optimization stage 2 (shared gradients and optimizer states). The precision was set to bfloat16 floating-point format. The parameters from the ‘model_5_multimer_v3’ preset defined by OpenFold were mostly carried over^46^. This preset has the largest MSA depths used during initial training and fine-tuning of the AlphaFold-Multimer parameters. We added the additional parameters from the AlphaFold finetuning preset: maximum additional MSA depth of 5120 sequences, and template module weight was set to 0.1. This was included in the multimer training preset to allow for fine-tuning during training. The full configuration file, and example training scripts are included in the supplementary information.

Each training epoch had 200 steps. The network was trained on eight NVIDIA L40S GPUs in a parallel format. The network was trained for 3 epochs total, with the checkpoint file saved as inference weights after 3 epochs. For the purposes of training on multimeric subunit assemblies, to better leverage the data, the Evoformer, MSA, and lDDT gradients were frozen during training. This resulted in only the structure module updating during training. The SID-ERMS data contains information about the interfaces in the protein complex, therefore we only wanted this information to be passed to the parts of the network that build the interfaces. This leverages the strengths of the AlphaFold Evoformer and MSA modules, while letting the SID-ERMS information construct higher accuracy interfaces between the protein subunits.

### Finetuning Inference on SID-ERMS data

For proteins with deep MSAs or very closely related templates, sequence- and template-derived features can often overshadow other sources of information during inference. For this reason, several deep-learning methods that incorporate experimental restraints artificially restrict MSA depth during structure prediction. Similar to Alphalink^22^ and DMSFold^69^, we sequentially subsampled each MSA to various effective depths (called N-effective, or Neff). For each MSA entry, the sequence similarity was calculated to the query sequence. Sequences with over 0.8 sequence similarity were rounded to 1.0. The MSA was then subsampled sequentially until the desired number of Neff sequences was reached. To ensure the predicted protein structures are in accordance with the experimental data, the main inference loop of the network was modified. Before each recycle, during inference, the intermediate structure was relaxed and the ERMS data was simulated. The RMSE was then calculated, and network recycling continued if the RMSE was greater than 0.15. Once the RMSE was less than 0.15, recycling iterations ended and the output structure was saved. A max recycle count of 100 was set to prevent indefinite recycling when the RMSE threshold could not be reached. This procedure differed during training, where simulation occurred only on the last iteration of the recycling. This variable recycling enabled iterative refinement of each prediction toward the target ERMS profile. Similar refinement strategies have been implemented in networks such as Distance-AF^36^. In the present study, this variable-recycling procedure improved the accuracy of protein-complex assembly during inference.

### SID-ERMS pLDDT score

To evaluate the predicted structures both in terms of model confidence and agreement with experimental data, we developed a hybrid confidence score. We used the average predicted local distance difference test (pLDDT) output for each SIDFold prediction, along with the RMSE of the structures’ simulated ERMS data to the experimental ERMS data. The two scores were combined to form the confidence score defined in Equation 4.

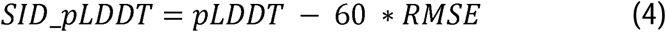

### RMSD, RMSD_100_ TM-score, and interface SASA comparisons

RMSDs were measured using PyMOL^74^. The Cα RMSD with no outlier rejection was measured for each predicted model against the reference structure. RMSD_100_^75^ measurements were calculated using the PyMOL RMSD, normalized to a 100 residue sequence length using Equation 5, where N is the total number of residues in the protein.

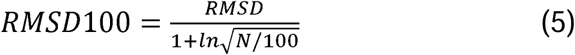

TM-scores were calculated using the US-align package^76^. Structures were aligned to the native structure using fully non-sequential alignment (-mm=5), all chains (ter=0), condensed output (outfmt=1).

Interface SASA’s were measured using PyRosetta’s InterfaceAnalyzer application^52^.

DockQ_i_ scores for multimeric structures were calculated using the DockQ v2 package^77^.

## Results and Discussion

### Integration of Experimental Data with an Auxiliary Loss Function Influenced Protein Structure

AlphaFold-Multimer prediction accuracy is notably lower than its monomer prediction accuracy. Most commonly, this inaccuracy is in the assembly of the subunits rather than the subunits themselves^49^. We hypothesized that guiding structure generation towards agreement with SID-nMS data would improve the accuracy of the predicted multimer structure. This was inspired by our previous work^44^, in which SID-ERMS was used to guide Rosetta docking. We further hypothesized that a deep-learning framework could learn relationships between the multivariate features of an ERMS profile and the structural organization of a protein complex, potentially improving upon our previous Rosetta-based approach. The SID-ERMS data contains information on the interface composition and connectivity for a protein complex. Therefore, we hypothesized that the AlphaFold structure module would be able to leverage this information to improve prediction accuracy. We thus implemented a range of functionalities into AlphaFold that facilitate SID-guided structure prediction. These functionalities were focused on the structure module, and include the ability to simulate SID-ERMS data as a core function of the AlphaFold structure generation process. The resulting network, termed SIDFold, is illustrated in Figure 1. The MSA-derived single and pair representations were generated using the standard AlphaFold architecture, while the structure-generation and recycling blocks were modified to incorporate SID-ERMS data. Unlike many experimentally guided AlphaFold-based methods, which embed experimental restraints directly into the residue level pair representation^22,36,37,78^, SIDFold uses a novel approach in which it incorporates experimental information through modifications to the structure module and recycling loop. The updated single and pair representations are input into the structure module, the recycling loop of which was modified to output an intermediate structure file during every recycle. Each intermediate structure was used to simulate an SID-ERMS profile using a PyRosetta implementation of the Rosetta `SID_ERMS_Prediction` application^53^, and the resulting profile was saved for the corresponding iteration. During training, this data was passed directly to the auxiliary SID-ERMS loss function, which evaluated the RMSE between the simulated and target ERMS profiles. During inference, the RMSE was calculated directly in the structure module to dynamically control the number of recycling iterations. Recycling terminated when the calculated RMSE fell below a predefined agreement threshold, at which point the output structure was saved. If the agreement threshold was not reached, the recycling loop continued until the threshold was met or the maximum number of recycles was reached. After each recycling iteration, the updated single and pair representations were returned to the Evoformer, and used as the inputs for the next recycling loop. Training metrics are presented in Supplementary Figure S1.

**Figure 1:**
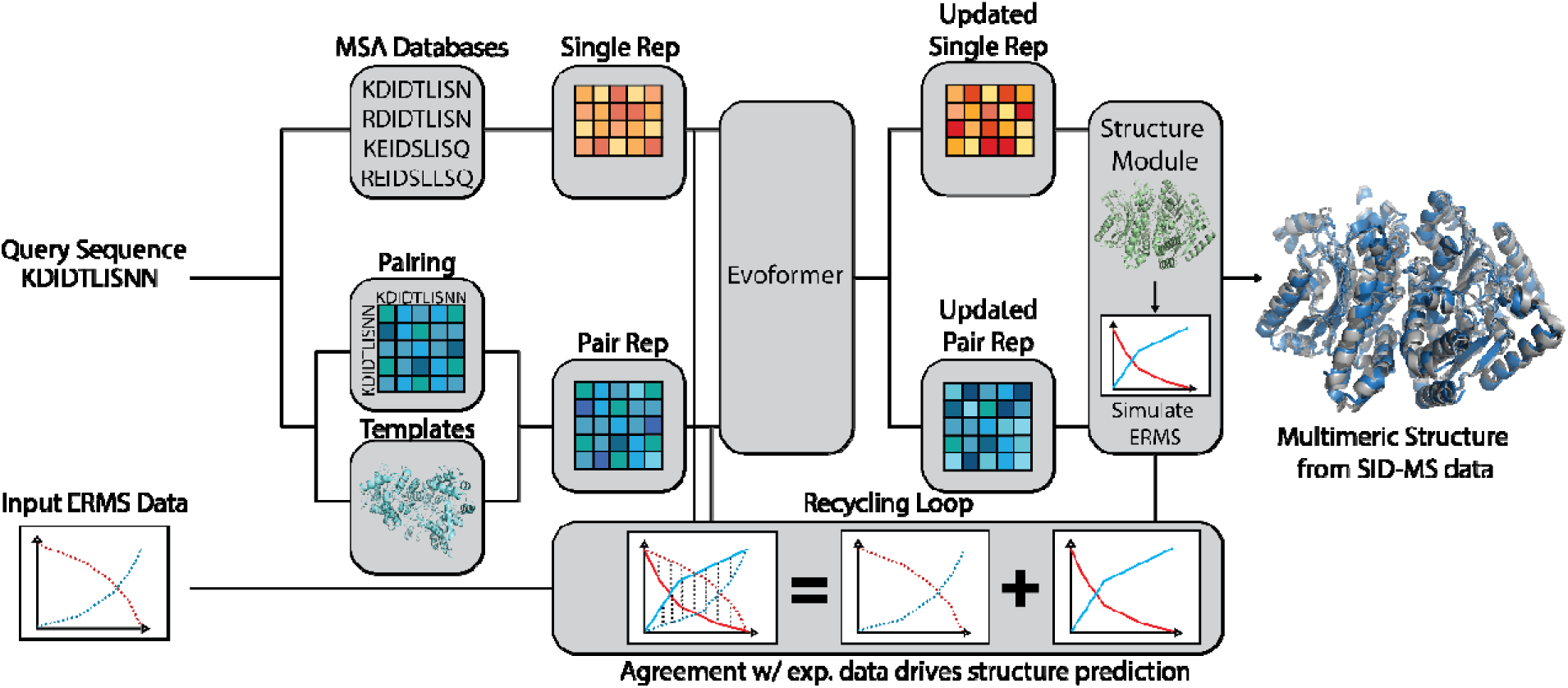
Network architecture for SIDFold. The SIDFold architecture was based on the AlphaFold2 architecture. The single and pair representations are initialized from the MSA and template databases. The representations are iterated and refined in the Evoformer. The updated representations are directly input to the structure module. Starting on the first cycle, the structure module outputs an intermediate structure every recycle. The SID-ERMS data for this intermediate structure is simulated, and during training this simulated data is input to the auxiliary SID-ERMS loss function (loss function not shown in graph). The loss function evaluates the difference between the experimental and simulated data to guide the network training. In inference, the simulated data is compared to the experimental data to guide the recycling loop. It will recycle the structure until a threshold level of data agreement is reached.

### SIDFold with simulated SID-ERMS data improved predictions over AlphaFold

We first benchmarked the network using simulated SID-ERMS data to evaluate its performance across a large and structurally diverse set of protein complexes. We used the BETA protein set, which is a dataset composed of protein structures released after AlphaFold was trained. We selected the multimeric proteins present in the dataset and applied the same structural filtering criteria used to construct the training dataset. This filtering was to enrich the benchmark set for complexes ideal for SID-nMS analysis and to prevent any data leakage from test structures being part of the AlphaFold training set. We simulated the SID-ERMS data for each protein to use as experimental input data into the network. For each benchmark target, five predictions were generated using SIDFold and AlphaFold, with the same five random seeds used across both methods. MSAs were subsampled to an N_eff_ of 10. This subsampling strategy was motivated by previous work demonstrating that reducing MSA depth can increase the influence of experimental restraints on the final prediction by limiting the otherwise dominant contribution of MSA-derived features during structure inference ^22,69^. We sought to simulate the average use case for which the native structure is not known prior to inference. To achieve this, first the SIDFold structures’ average pLDDT were reweighted using the SID_pLDDT metric (see Methods). Afterwards, for both AlphaFold and SIDFold, for each set of five seeded predictions, all models with an average pLDDT/SID_pLDDT > 70 (high confidence backbone models) had their RMSD averaged. If none of the 5 models had a pLDDT > 70, the highest average pLDDT/SID_pLDDT structure was used. The benchmark set accuracy (as measured by model RMSD) compared to AlphaFold can be seen in Figure 2A. We saw increased accuracy (i.e., decreased RMSD) for 138 of 227 structures when using SIDFold as compared to AlphaFold. For those that improved, the average improvement in accuracy was 3.42 Å RMSD. Furthermore, 93 of 227 proteins showcased an RMSD under 5 Å, which is a highly accurately predicted complex. We saw 27 of 227 proteins increase in accuracy from above 5 Å RMSD with AlphaFold to below 5 Å RMSD with SIDFold. This constituted a significant increase in accuracy, especially considering that only 7 predictions worsened in accuracy from below 5 Å with AlphaFold to above 5 Å RMSD with SIDFold. Therefore, SIDFold showed a consistent increase in accuracy across the dataset in comparison to AlphaFold.

**Figure 2:**
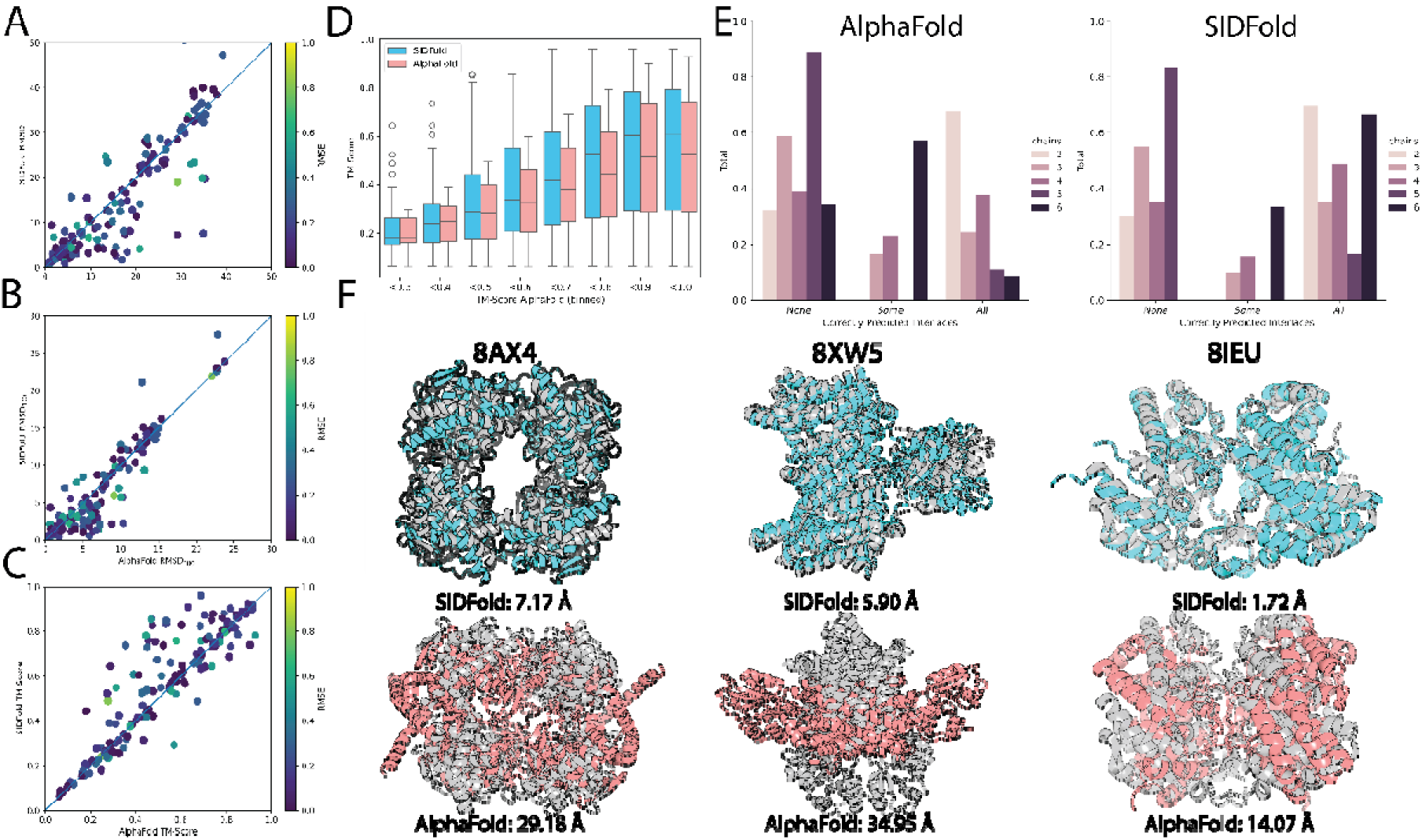
SIDFold predictions with simulated data. Five seeded inference predictions were performed for both AlphaFold and SIDFold. (A) RMSD comparison between SIDFold and AlphaFold predictions for 227 proteins outside of AlphaFold’s training set with simulated SID-ERMS data. 138/227 structures improved in raw RMSD, with an average improvement of 3.42 Å for proteins that improved. 93/227 structures showed a complex RMSD below 5 Å, compared to only 73/227 AlphaFold predicted structures having an RMSD below 5 Å. Colors of the points denote the average RMSE of the simulated SID-ERMS data of the predicted SIDFold models. (B) RMSD_100_ comparison between SIDFold and AlphaFold predictions for 227 proteins outside of AlphaFold’s training set with simulated SID-ERMS data. 139/227 structures improved in raw RMSD_100_, with an average improvement of 1.84 Å for proteins that improved. 138/227 structures showed a complex RMSD_100_ below 5 Å, compared to 111/227 AlphaFold predicted structures having an RMSD_100_ below 5 Å. Colors of the points denote the average RMSE of the simulated SID-ERMS data of the predicted SIDFold models. (C) Sequence independent TM-score comparisons between SIDFold and AlphaFold predictions for the 227 proteins outside of AlphaFold’s training set with simulated SID-ERMS data. 140/227 structures improved in raw TM-score, with an average improvement of 0.08 TM-score for proteins that improve. 55/227 SIDFold predictions showed a TM-score higher than 0.8, compared to 41/227 AlphaFold predictions. (D) TM-Score distributions of AlphaFold (pink) and SIDFold (cyan), binned to TM-Scores of AlphaFold predictions. (E) DockQ_i_ analysis for SIDFold and AlphaFold predictions on the 227 proteins in the benchmark dataset. Inaccurate interfaces were defined as DockQ_i_ < 0.23. Averaged across all oligomeric states, AlphaFold had a 49.4% accuracy for interface prediction. SIDFold had an accuracy of 59.3% for interface prediction. (F) SIDFold predictions and AlphaFold predictions subsampled to Neff=10 overlayed with the native structure for three proteins from the benchmark dataset. For both SIDFold and AlphaFold, structures with the individual RMSD closest to the average reported in Figure 2A are shown. AlphaFold predictions are colored pink, SIDFold predictions are labeled cyan, and native structures are colored gray. RMSDs of the individual structures are reported below the aligned structures.

Because RMSD is a size dependent measurement, comparisons between larger complexes can yield disproportionately large and potentially misleading values^75^. Therefore, we also evaluated model accuracy using RMSD_100_, which is a size-independent RMSD normalized to a sequence length of 100 residues. The RMSD_100_ comparison between SIDFold and AlphaFold is shown in Figure 2B. Similar to the results shown in Figure 2A, SIDFold produced lower RMSD_100_ values for 139 of the 227 complexes in the benchmark set. Additionally, 138 of 227 SIDFold predictions achieved an RMSD_100_ below 5 Å with 20 complexes significantly increasing in accuracy, defined as having above 5 Å RMSD_100_ when predicted with AlphaFold and below 5 Å RMSD_100_ when predicted with SIDFold. Both RMSD and RMSD_100_ are sequence dependent alignments, potentially artificially increasing the reported accuracy when comparing a prediction to its native structure. For this reason, we also performed a sequence independent TM-score alignment to probe the accuracy of the predictions without any guidance from a sequence alignment. The results of this TM-score comparison can be seen in Figure 2C. We observed similar trends to the RMSD and RMSD_100_, where 140 of 227 complexes showed an increase in accuracy with SIDFold compared to AlphaFold. In total, 55 SIDFold predictions achieved a TM-score of 0.8 or greater. For 20 complexes, SIDFold increased the TM-score from below 0.8 with AlphaFold to above 0.8. Together, these analyses indicate that the improvement in accuracy obtained with SIDFold was retained after accounting for both complex size and the potential influence of sequence-guided structural alignments. Furthermore, we binned the TM-Score predictions for both networks to the AlphaFold TM-Scores to directly compare the TM-Scores at all accuracies (Figure 2D). Across all ranges of AlphaFold accuracies, SIDFold consistently outperformed AlphaFold.

We then endeavored to test the accuracy of the predicted interfaces present in the structures. We directly compared SIDFold against AlphaFold using DockQ_i_, the results of which can be seen in Figure 2E. An interface was defined as correct if it had a DockQ_i_ > 0.23, following established benchmarking protocols^49^. Predicted structures were then binned into three categories: ‘No interfaces predicted correctly’, ‘Some interfaces predicted correctly’, and ‘All interfaces predicted correctly’. Averaging across all oligomeric states, AlphaFold’s multimeric accuracy was 49.4%, defined as some or all interfaces predicted correctly. This matched previous results (ref ^49^) almost exactly, showing our benchmark set was a representative sample for measuring the performance of AlphaFold. Additionally, AlphaFold’s accuracy worsened with higher oligomeric state proteins, matching the previous reported findings. Counting the structures with all correct interfaces, AlphaFold’s accuracy for dimers was 67.5%, 24.5% for trimers, 37.7% for tetramers, 11.1% for pentamers, and 8.6% for hexamers. SIDFold’s multimeric accuracy averaged across all oligomeric states was 59.3%. Counting the structures with all correct interfaces, SIDFold’s accuracy for dimers was 69.7%, 45.0% for trimers, 64.7% for tetramers, 16.6% for pentamers, and 66.7% for hexamers. This was notably higher than AlphaFold’s accuracy, across every single category, showing a significant improvement over the network’s accuracy. Additionally, SIDFold did not decrease in accuracy with higher oligomeric states. On the contrary, when isolating the hexameric predictions, SIDFold predicted all hexameric interfaces correctly in 66.7% of cases. In comparison, AlphaFold only predicted 8.6% of the hexameric predictions completely correctly. This is a significant increase in the accuracy of higher oligomeric state complexes and underscores that SID-nMS data is useful in guiding interface prediction. Notably, SIDFold outcompeted AlphaFold at every oligomeric state in accuracy (see Figure 2E), and did not degrade in accuracy with higher order complexes.

Three candidate structural alignments can be seen in Figure 2F. Native structures are colored gray, SIDFold structures are colored cyan, AlphaFold predictions are colored pink. The proteins shown are membrane-bound PRTase (PDB ID: 8J8J), *B. bacteriovorus* secreted protein Bd1399 (PDB ID: 8OKH), and DUF2891 family protein CJ0554 (PDB ID: 8IEU). Each protein’s RMSD decreased by over 10 Å with SIDFold compared to AlphaFold. Additionally, AlphaFold failed to predict the interface of each complex. SIDFold restored the native interface structure, highlighting the power of integrating multimeric structure data for increasing multimeric prediction accuracy.

### SIDFold with experimental SID-ERMS data improved predictions over AlphaFold

After benchmarking SIDFold on simulated data, we tested the network on an experimental dataset. The experimental benchmark set contained 20 protein complexes with high-quality SID-ERMS data and high-resolution crystal structures. The dataset consisted of six dimers, ten tetramers, three pentamers, and one hexamer. PDB ID’s and point groups for each complex can be found in Supplementary Table S2. These proteins provided a diverse set of oligomeric states to evaluate our model. For comparison we ran five seeded predictions (using the same five seeds across the entire experimental benchmark set) using AlphaFold subsampled to a Neff 10 for each protein in our experimental set. To identify high confidence predicted models, we applied the SID_pLDDT to each SIDFold prediction using the models’ average pLDDT and the SID-ERMS data (see Methods). The comparison of structural accuracy (as measured by model RMSD) between AlphaFold and SIDFold predictions can be seen in Figure 3A.

**Figure 3:**
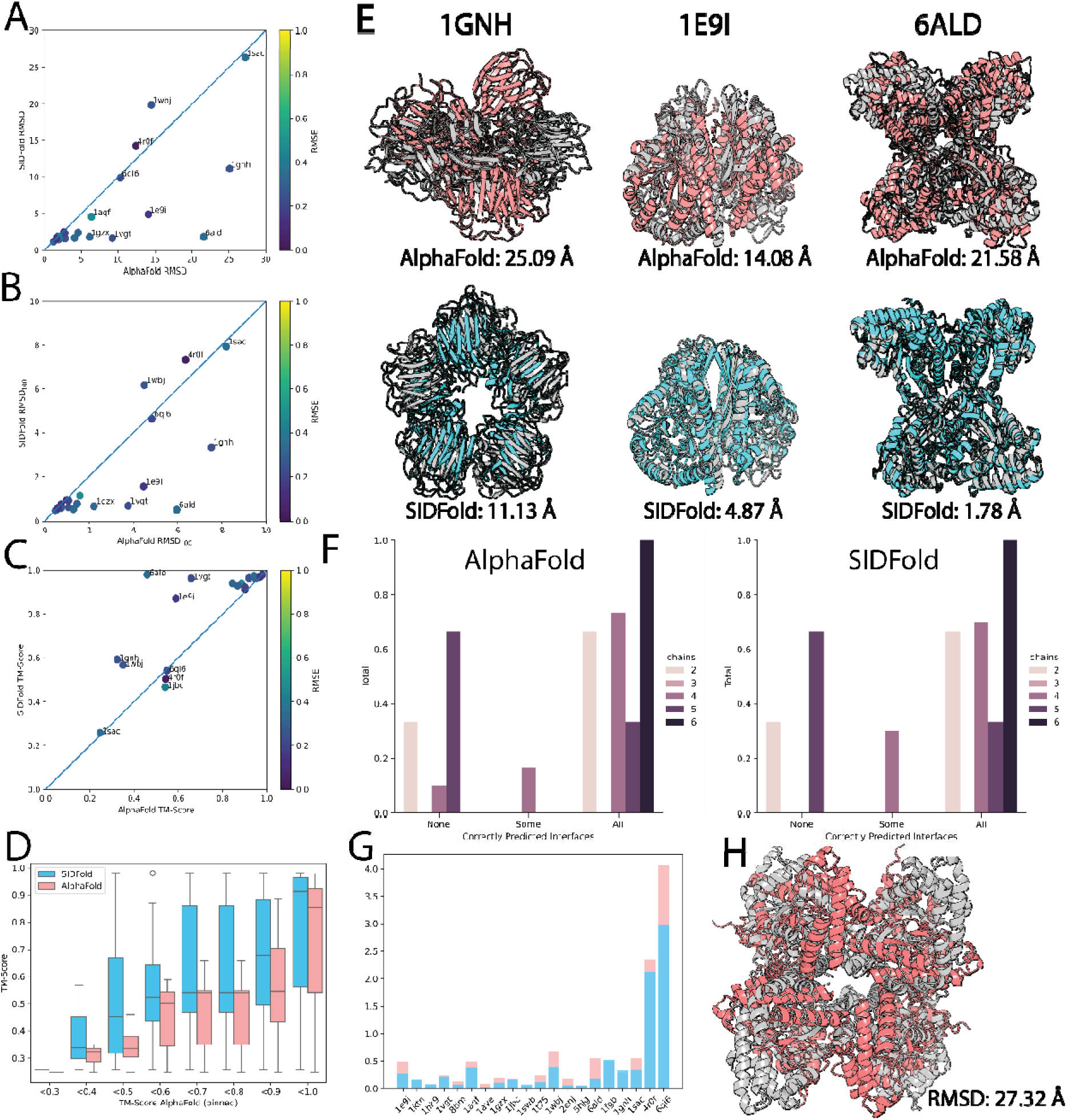
SIDFold predictions with experimental data. Five seeded inference predictions were performed for both AlphaFold and SIDFold. (A) RMSD comparison between SIDFold and AlphaFold predictions for 20 proteins with experimental SID-ERMS data. 18/20 structures improved in raw RMSD, and 15/20 structures showed a complex RMSD below 5 Å, compared to only 10/20 AlphaFold predicted structures having an RMSD below 5 Å. Colors of the points denote the average RMSE of the simulated SID-ERMS data of the predicted SIDFold models. (B) RMSD_100_ comparison between SIDFold and AlphaFold predictions for 20 proteins with experimental SID-ERMS data. 18/20 structures improved in raw RMSD_100_, 15/20 structures showed a complex RMSD_100_ below 2 Å, compared to 11/20 AlphaFold structures. Colors of the points denote the average RMSE of the simulated SID-ERMS data of the predicted SIDFold models. (C) Sequence independent TM-score comparisons between SIDFold and AlphaFold predictions for the 20 proteins in the experimental set. 17/20 proteins improved in TM-score, and 14/20 SIDFold predictions showed a TM-score higher than 0.8, compared to 11/20 AlphaFold predictions. (D) TM-Score distributions of AlphaFold (pink) and SIDFold (cyan), binned to TM-Scores of AlphaFold predictions. (E) SIDFold predictions and AlphaFold predictions overlayed with the native structure for three proteins from the experimental dataset. For both SIDFold and AlphaFold, structures with the individual RMSD closest to the average reported in Figure 3A are shown. AlphaFold predictions are colored pink, SIDFold predictions are labeled cyan, and native structures are colored gray. RMSDs of the individual structures are reported below the aligned structures. (F) DockQ_i_ analysis for SIDFold and AlphaFold predictions on 20 proteins in the experimental set. Inaccurate interfaces were defined as DockQ_i_ < 0.23. Averaged across all oligomeric states, AlphaFold had a 72.7% accuracy for interface prediction. SIDFold had an accuracy of 75.0% for interface prediction. (G) Interface SASA measurements for SIDFold predictions versus AlphaFold predictions for the 20 proteins in the experimental set. Interface SASA calculated using Rosetta InterfaceAnalyzer in PyRosetta. The SASA for the predicted structures and native structures are measured, and the difference between the two measurements normalized to the native structure are shown. A perfect match in interface SASA would be 0.0, and a 100% error relative to the native interface would be 1.0. AlphaFold predictions are shown in pink, SIDFold predictions are overlayed in cyan. 14/20 proteins in the evaluation set had a lower error in interface SASA when comparing SIDFold to AlphaFold. (H) SIDFold inference for 6ALD using nonsense data. The nonsense prediction is colored red, and the native structure is labeled gray. The RMSD is reported underneath the aligned structures.

For 18 of the 20 complexes in our evaluation set, we saw improvement in RMSD for SIDFold predictions in comparison to the AlphaFold predictions. Additionally, 15/20 proteins predicted with SIDFold had an RMSD below 5 Å, denoting a highly accurate protein complex structure. This was in comparison to AlphaFold predictions, in which only 10/20 predictions were below 5 Å RMSD. Hence, of the 18/20 proteins that improved in RMSD, 5 predictions improved from above 5 Å RMSD with AlphaFold to below 5 Å RMSD with SIDFold. The protein D-sialic acid aldolase (PDB ID: 6ALD) exhibited an RMSD improvement of 20 Å (see Figure 3A). Furthermore, the only two proteins in the set that worsened were inaccurate structures (AlphaFold RMSD > 12 Å), and they only worsened by 2 Å and 5 Å RMSD, respectively. The RMSD_100_ comparison for the same experimental benchmark set of predictions can be seen in Figure 3B. A similar trend to the RMSD comparison was observed, where 18 of 20 proteins improved in RMSD_100_ for SIDFold predictions compared to AlphaFold. 15/20 structures had an RMSD_100_ below 2 Å, compared to 11/20 Alphafold predictions. This served as further verification of the robustness of SIDFold’s improved performance compared to AlphaFold, showing improved accuracy independent of protein size.

We additionally performed a sequence independent TM-score analysis. The results of the TM-score analysis can be seen in Figure 3C. Similar to the results of the RMSD and RMSD_100_ analysis, 17/20 proteins improved in TM-score with SIDFold predictions compared to AlphaFold. Furthermore, 14/20 SIDFold predictions had a TM-score above 0.8, indicating highly accurate structures, whereas only 11/20 structures predicted with AlphaFold reached this threshold. 5/20 AlphaFold predictions had a TM-score below 0.5, indicating an incorrect overall structure, compared to only 2/20 SIDFold predictions with a TM-score below 0.5. Furthermore, we again binned the TM-Score distributions for both networks to AlphaFold’s TM-Score, to compare the performance across all accuracies (Figure 3D). As with the simulated data, across all ranges of AlphaFold accuracies, SIDFold consistently outperformed AlphaFold. For a vast majority of the evaluation set, across multiple analysis metrics, SIDFold gave a nearly unanimous, and in several cases a drastic, improvement over un-guided deep learning methods. Overall, this was a highly significant improvement over the purely computational AlphaFold method, highlighting the power of integration of experimental data into deep learning networks.

Structural alignments of three proteins in the experimental set are shown in Figure 3E. Native structures are colored gray, SIDFold structures are colored cyan, AlphaFold predictions are colored pink. The proteins shown are: C-reactive protein, a C5 pentamer (PDB ID: 1GNH), Enolase, a C2 dimer (PDB ID: 1E9I), and D-sialic acid aldolase, a D2 tetramer (PDB ID: 6ALD). These three proteins showcase the structural and oligomeric diversity present in the experimental set. For all three proteins, AlphaFold incorrectly predicted the interfaces, and thus the multimeric structure. In the case of the pentamer, 1GNH, AlphaFold furthermore predicted an incorrect molecular geometry, adopting a square pyramidal geometry instead of a pentagonal planar geometry. SIDFold restored the correct interface for all three structures. Additionally, SIDFold restored the correct molecular geometry for the C5 pentamer. This highlights the value of the interface information encoded in SID-ERMS data.

We also performed a DockQ_i_ interface analysis for the experimental set. Again, an interface was defined as correct if it had a DockQ_i_ > 0.23. Predicted structures were binned into three categories: ‘No interfaces predicted correctly’, ‘Some interfaces predicted correctly, and ‘All interfaces predicted correctly’. Averaged across all oligomeric states, AlphaFold had a multimeric structure accuracy of 72.5%, while SIDFold had an accuracy of 75% (see Figure 3F). These accuracies were much higher than the values reported for the benchmark dataset above. The likely reason for this is that the experimental dataset contained very well studied, characterized, and ordered proteins. This likely resulted in a higher baseline accuracy for these proteins compared to the typical protein in the benchmark set. However, SIDFold still exhibited a higher accuracy than AlphaFold, such as predicting 100% of the tetramers accurately. This shows that even in ideal cases like the proteins in the experimental set, SIDFold outperformed AlphaFold in multimeric accuracy.

To further validate whether the network is integrating the SID-ERMS data directly, we performed two additional analyses. First, for the final evaluation set of experimental structures, we compared the interface Solvent-Accessible-Surface-Area (interface SASA), for each seeded predicted SIDFold and AlphaFold structure. The interface SASA is directly used to calculate the SID-ERMS data, making it the ideal structural metric to use to validate the integration of SID-ERMS data into the network. The results of this comparison can be seen in Figure 3G. The interface SASA for each structure was measured using Rosetta’s `InterfaceAnalyzer` in PyRosetta. First, the native structure’s interface SASA was measured. Then, the predicted structure’s interface SASA was measured, and the difference between the two measurements was normalized to the native interface SASA (data shown in Figure 3G for SIDFold and AlphaFold). If the predicted structure’s interface SASA perfectly matched the native structures, the normalized difference would be 0.0. A 100% difference in interface SASA would be 1.0. AlphaFold predictions are colored pink, and SIDFold predictions are overlayed in cyan. In 14/20 cases, we saw improvement in interface SASA relative to AlphaFold, with an average improvement of 11.5% across the 20 proteins. Furthermore, in the 6/20 cases we did not improve, the average difference between SIDFold and AlphaFold was less than 0.1 (10%). Therefore, SIDFold was able to match or improve the accuracy of the interface SASA across almost all predictions in the set. Furthermore, the two proteins with the highest error, Lysozyme (PDB ID: 4R0F) and Beta-lactoglobulin (PDB ID: 6QI6), are proteins that were incorrectly predicted in both AlphaFold and SIDFold (minimum RMSD for all predicted models > 10 Å). This further supports that the network is directly integrating the SID-ERMS data into its structural prediction calculations.

Our second analysis used nonsense SID-ERMS data for protein inference. Our hypothesis was that if the network is using the SID-ERMS data for the structure prediction, feeding nonsense data would result in an inaccurate prediction. We chose the D2 tetramer D-sialic acid aldolase (PDB ID: 6ALD) as a test system, as that protein showed a high increase in accuracy with SIDFold relative to AlphaFold. To create the nonsense data, we set every frequency value in the experimental data with 0. This would indicate no intact or fragmented complexes are present, which is not a physically achievable result in SID-nMS. The results of this inference can be seen in Figure 3H. The native structure is colored gray, and the nonsense SIDFold prediction is colored red. The nonsense data drove the predicted structure into a C4 symmetry instead of the native D2 symmetry, resulting in a completely incorrect multimeric structure (RMSD = 27.32 Å). This provided strong evidence that the network is directly using the SID-ERMS data in the structure prediction, as incorrect data resulted in an inaccurate prediction.

### SIDFold predictions outperformed current state-of-the-art Rosetta SID-guided modeling

Rosetta rescoring of docked models with SID-ERMS data is the current leading method for integrating SID-nMS data into protein structure prediction^44^. With SIDFold being the first method to use machine learning and SID-nMS data, we aimed to compare its performance to `SID_ERMS_Rescore` in Rosetta. In contrast to the 20 complete protein complex models generated with SIDFold (Figure 3), `SID_ERMS_Rescore` was able to only predict the dimeric form of the three C5 pentamers (PDB IDs: 1SAC, 1GNH, 1FGB) and C6 hexamer (PDB ID: 1HK9) in our experimental set, preventing a direct comparison between the two methods for these four proteins. The comparison between the 16 structures for which we predicted full complex structures with both methods is shown in Figure 4. For the SIDFold structures, we used the same five seeded predictions with regraded SID_pLDDT scores seen in Figure 3. For the SID_ERMS_Rescore in Rosetta structures, we identified the top scoring model of 10,000 docked structures from ^44^. 13/16 proteins were predicted below 5 Å RMSD with respect to the native structure with SIDFold. In comparison, SID_ERMS_Rescore predicted only 3/16 proteins (PDB IDs: 1JBC, 1T75, 5HJG) under 5 Å RMSD with respect to the native. This constitutes a significant improvement in accuracy, again showcasing the power of deep learning methods with SID-nMS data. Furthermore, the improvements of SIDFold go beyond the accuracy of the structures. SID_ERMS_Rescore is reliant on an accurate monomeric structure, which is then docked and rescored using SID-ERMS data. This constitutes a limit on its general use cases, restricting application to cases where the monomeric structure is known (or can be accurately predicted) but the multimeric structure is not. This contrasts with SIDFold, which constitutes an entirely *de novo* process requiring no prior knowledge of the structure. The prior knowledge of the monomer structure was what allowed `SID_ERMS_Rescore` to produce more accurate models in 3/16 proteins (PDB IDs: 4R0F, 6QI6, 1WBJ) in the set over SIDFold. Two of the three benchmark proteins were dimers (PDB IDs: 4R0F, 6QI6) and started with the correct monomer structure for docking. This is why the predicted complexes were more accurate than those from SIDFold, which generated the monomeric structure without prior knowledge before complex assembly. However, for all three predictions, neither method ultimately produced a high accuracy complex (RMSD < 5 Å), and the average RMSD improvement across all 16 proteins was 8.5 Å, underscoring how significantly SIDFold outperformed the current leading SID-guided prediction method.

**Figure 4:**
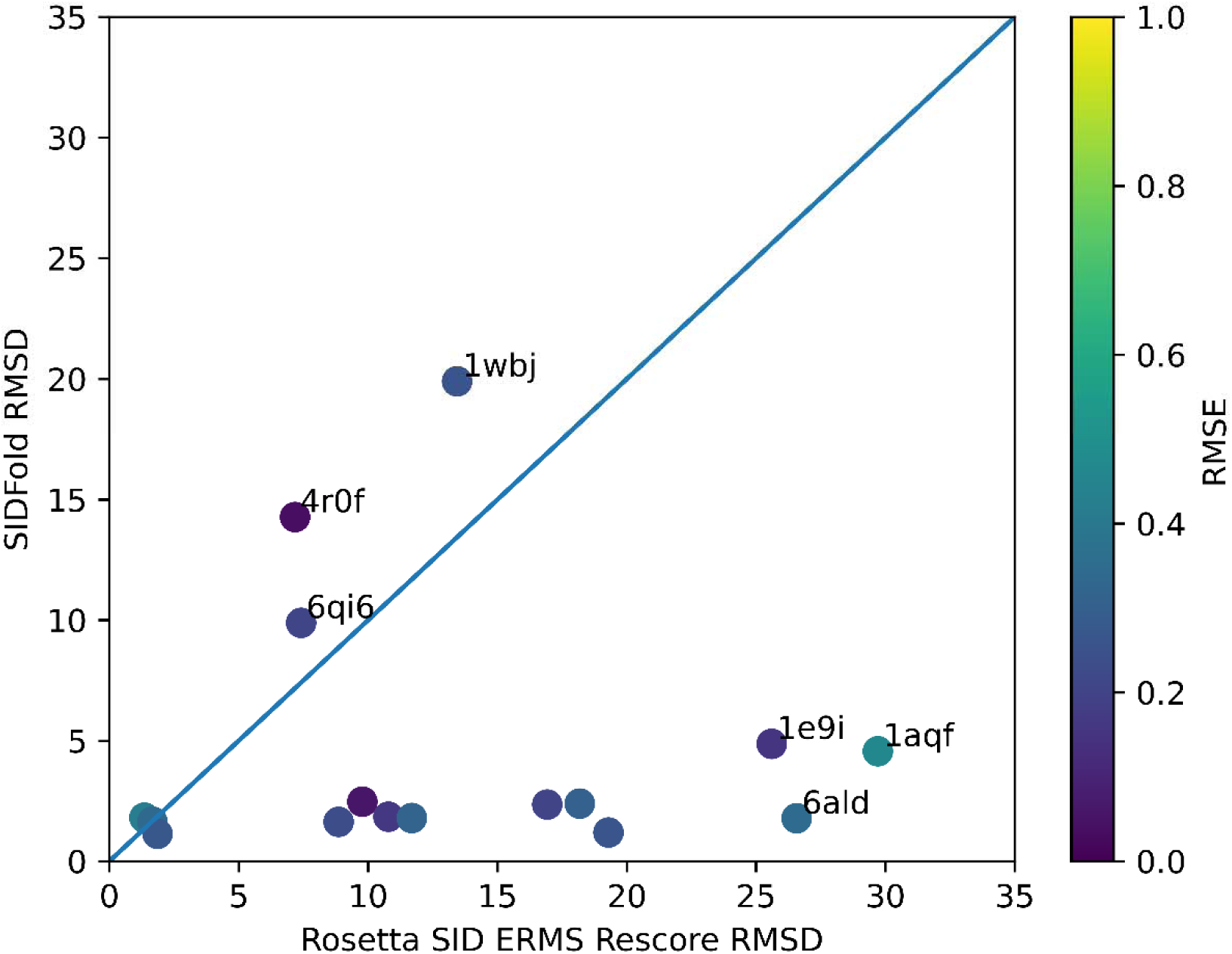
SIDFold predictions compared to SID ERMS Rescore in Rosetta. Comparison of the 16 structures fully predicted with both SIDFold and SID ERMS Rescore in Rosetta. SIDFold structures are the same 5-seeded predictions shown in Figure 3. SID ERMS Rescore structures are the top scoring models from 10,000 docked structures, respectively. Colors of the points show the average RMSE of the simulated SID-ERMS data of the predicted SIDFold models. 13/16 structures predicted with SIDFold had less than 5 Å to the native structure, compared to only 3/16 with SID ERMS Rescore.

## Conclusion

SID-nMS is a versatile, adaptable, and rapid technique that provides information on the stoichiometry and connectivity of protein complexes. Previous work developed a docking-based method for integrating SID-nMS data into multimer structure prediction. However, inherent limitations remained and no machine-learning based method existed that integrates SID-nMS into its network. In this work, we present the first of its kind deep-learning network that incorporates SID-nMS data, and the first deep-learning network to incorporate any kind of native MS data. The network, SIDFold, is derived from the AlphaFold network, with several augmentations. We developed a novel auxiliary loss function and recycling loop that uses SID-ERMS data to guide the structural assembly of the predicted complex. We benchmarked this network using proteins that were deposited into the PDB after AlphaFold was trained. We simulated the SID-ERMS data for each protein using the native structure. This simulated data was used to predict the protein complex, and compared with AlphaFold. We saw increased accuracy in 138/227 complexes, with no significant decrease in accuracy for the other complexes in the set. We then performed the same comparison using 20 proteins with experimental SID-ERMS data as a final evaluation. We saw 18/20 proteins increased in accuracy, with one protein decreasing its RMSD by 20 Å. Additionally, SIDFold accurately predicted the correct interfaces in several cases where AlphaFold incorrectly assembled the multimer protein. Finally, we compared SIDFold to the previously developed SID_ERMS_Rescore Rosetta method. SIDFold showed higher accuracy for 13/16 proteins compared to SID_ERMS_Rescore.

The SIDFold network is the first of its kind to integrate native MS, and more specifically SID-nMS, for multimeric structure determination. This method outperforms AlphaFold and the previously published Rosetta docking method in structural accuracy, including predictions of more accurate interfaces. The combination of native MS and deep learning proves to be powerful in light of the growing prevalence of nMS and the ability of deep learning to convert the data into structure. This network is freely available on the Lindert lab github page (https://github.com/LindertLab/SIDFold). Additionally, example config files and command line inputs can be found in the Supplementary Information. While this network significantly outperforms other published methods, there are potential avenues for improving SIDFold in future work. Higher level embeddings such as Multi-Layer Perceptrons (MLPs) or Locally Linear Embeddings (LLEs) offer potential to decode higher complexity relationships in the SID-ERMS data. Additionally, other methods of data simulation like Molecular Dynamics simulations can give more structural sampling to mimic higher charge states. These serve as potential next steps to improve the accuracy of the predicted structure in conjunction with experimental SID-nMS data.

## Acknowledgements

We would like to thank fellow Lindert lab members for their insight and conversations regarding this project. We additionally thank the Wysocki lab for their input on simulating the SID-ERMS data. This work used computational and storage services associated with the Hoffman2 Cluster which is operated by the UCLA Office of Advanced Research Computing’s Research Technology Group. This work was supported by NIH (RM1-GM149374, to V.H.W. and S.L.).

