## Supplemental Information for "Incorporating Surfaced-Induced Dissociation Mass Spectrometry Data into an AlphaFold-derived deep learning network improves protein structure prediction"

**Supplementary Information**

| Point group | C2 Dimer | C3 Trimer | C2 Tetramer | D2 Tetramer | C1 Pentamer |
| --- | --- | --- | --- | --- | --- |
| File Contents | A B  A_B | A B C  A_B B_C C_A | A B C D  A_B C_D  B_C | A B C D  A_B C_D  A_D B_C  A_C B_D | A B C D E  A_B  A_C  A_D  A_E  B_C  B_D  B_E  C_D  C_E  D_E |

**Table S1: Example connectivity files for various point groups.** The table contains the point group and oligomeric state for an example protein complex, and additionally the file contents in a line-by-line format for the connectivity file which encodes the point group and oligomeric state for SID-ERMS data simulation.

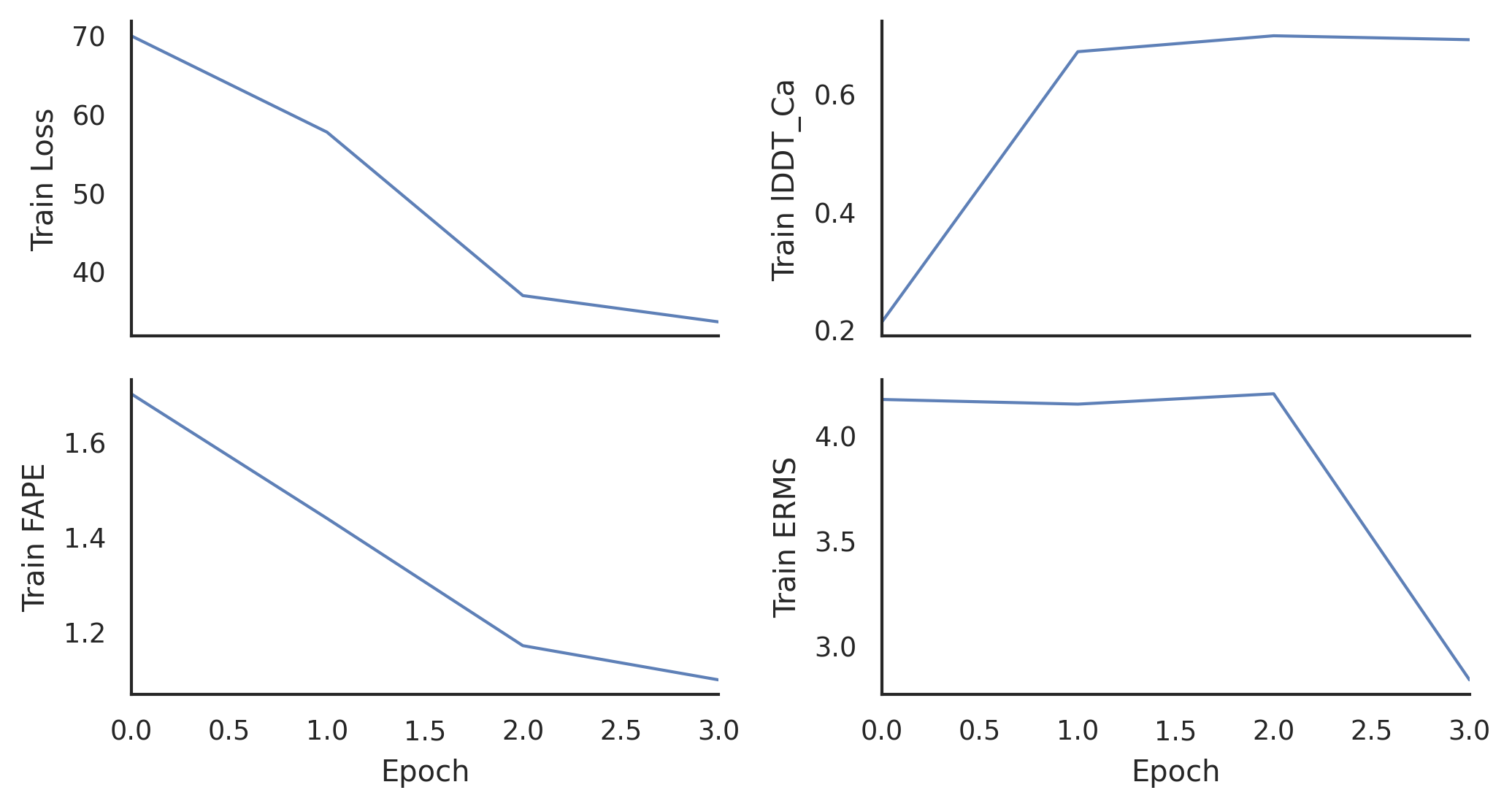

**Figure S1: Training metrics for SIDFold network.** Each epoch had 200 steps. The training was spread across 8 L40S GPUs, and the network trained for 3 epochs. (A) The total loss per epoch during training. (B) The local distance difference test for Cα atoms (lddt_ca) loss per epoch during training. (C) The frame aligned point error (fape) loss per epoch during training. (D) The auxiliary ERMS loss per epoch during training.

| Name | PDB ID | Complex Type | Symmetry |
| --- | --- | --- | --- |
| Enolase | 1E9I | homodimer | C2 |
| DeoC | 1KTN | homodimer |  |
| IspD | 1VGT | homodimer |  |
| Lysozyme | 4R0F | homodimer |  |
| Beta-lactoglobulin | 6QI6 | homodimer |  |
| Triose phosphate isomerase | 8TIM | homodimer |  |
| Pyruvate kinase | 1AQF | homotetramer | D2 |
| Avidin (neutravidin used experimentally) | 1AVE | homotetramer |  |
| Concanavalin A | 1JBC | homotetramer |  |
| Streptavidin | 1SWB | homotetramer |  |
| Carbonic anhydrase | 1T75 | homotetramer |  |
| Uracil phosphoribosyl transferase | 2EHJ | homotetramer |  |
| Transthyretin | 5HJG | homotetramer |  |
| D-sialic acid aldolase | 6ALD | homotetramer |  |
| Hemoglobin | 1GZX | heterotetramer | Pseudo-D2 |
| Tryptophan synthase | 1WBJ | heterotetramer | C2 |
| Cholera toxin B | 1FGB | homopentamer | C5 |
| C-reactive protein | 1GNH | homopentamer |  |
| Serum amyloid P | 1SAC | homopentamer |  |
| Hfq | 1HK9 | homohexamer | C6 |

**Table S2: The 20 proteins comprising the experimental evaluation dataset.** Information is presented on the name, PDB ID, complex type, and symmetry point group for each protein complex.

**Code for connectivity file generation**

To run the Quatsymm java executable to obtain the symmetry point group for a given protein:

java -Xmx2G -jar "<path to quatsymm directory>/quatsymm-2.3.0.jar" "$@"

The following is an example output for one protein:

| Name | Size | Subunits | Stoichiometry | Pseudostoichiometry | Symmetry | Local | Method | SymmRMSD | SymmTMscore |
| --- | --- | --- | --- | --- | --- | --- | --- | --- | --- |
| 7n8r | 2 | [A,B] | A2 | false | C2 | false | C2_ROTATION | 1.32 | 0.04 |

Using this symmetry, an example code segment (in python) for generating a connectivity file for this point group is:

for chain in chain_list: with open(”output.csv", 'a+') as file: file.write(chain[0]+' ')	with open(“output.csv", 'a+') as file: file.write('\n')	file.write(chain_list[0]+'_'+chain_list[1])

**Tutorial for running SIDFold**

Example command for running inference with SIDFold:

<path to conda env>/bin/torchrun \

--nnodes=$NNODES \

--nproc_per_node=$NPROC_PER_NODE \

--node_rank=$PMI_RANK \

--rdzv_id=manualtest$JOB_ID \

--rdzv_backend=c10d \

--rdzv_endpoint=${MASTER_ADDR}:${MASTER_PORT} \

<path to SIDFold dir>/SIDFold/run_pretrained_openfold.py \

<path to fasta dir> \

<path to databases>/mmcif_files \

--use_precomputed_alignments <path to MSA alignments> \

--output_dir <path to output dir> \

--uniref90_database_path <path to databases>/uniref90.fasta \

--mgnify_database_path <path to databases>/mgnify/mgy_clusters_2022_05.fa \

--pdb70_database_path <path to databases>/pdb70 \

--uniclust30_database_path <path to databases>/uniclust30_2018_08 \

--kalign_binary_path <path to binaries>/kalign \

--bfd_database_path <path to databases>/bfd_metaclust_clu_complete_id30_c90_final_seq.sorted_opt \

--hhblits_binary_path <path to binaries>/hhblits \

--hhsearch_binary_path <path to binaries>/hhsearch \

--jackhmmer_binary_path <path to binaries>/jackhmmer \

--neff 10 \

--preset reduced_dbs \

--config_preset "model_5_multimer_v3" \

--model_device "cuda" \

--jax_param_path <path to weights>/model_5_multimer_v3.npz

**Example command used to train SIDFold**

<path to conda env>/bin/torchrun \

--nnodes=$NNODES \

--nproc_per_node=$NPROC_PER_NODE \

--node_rank=$PMI_RANK \

--rdzv_id=manualtest$JOB_ID \

--rdzv_backend=c10d \

--rdzv_endpoint=${MASTER_ADDR}:${MASTER_PORT} \

<path to SIDFold dir>/SIDFold/train_openfold.py \

<path to train dir>/mmcif_train/ \

<path to alignments database dir>/ \

<path to mmcif database>/mmcif_files \

<path to output dir>/ \

{template date cutoff} \

--train_mmcif_data_cache_path <path to training dir>/train_mmcif_cache.json \

--val_mmcif_data_cache_path <path to training dir>/val_mmcif_cache.json \

--val_data_dir <path to training dir>/mmcif_val/ \

--val_alignment_dir <path to training dir>/val_alignments/ \

--kalign_binary_path <path to binaries>/kalign \

--template_release_dates_cache_path <path to training dir>/template_release_cache.json \

--deepspeed_config_path <path to training dir>/deepspeed_config.json \

--obsolete_pdbs_file_path <path to databases>/obsolete.dat \

--use_small_bfd True \

--seed <random int> \

--gpus <num gpus> \

--num_nodes 1 \

--checkpoint_every_epoch \

--train_epoch_len <integer> \

--max_epochs <integer> \

--config_preset "model_5_multimer_v3" \

--resume_from_jax_params <path to weights>/params_model_5_multimer_v3.npz \

--precision bf16-mixed \

--alignment_index_path <path to alignment index>/super.index

**Supplementary Dataset 1**

SIDFold_Supp.tar.gz contains:

1. training_pdbs.csv: a csv file containing the PDB ID’s and oligomeric state of all 200 proteins in our training dataset
2. validation_pdbs.csv: a csv file containing the PDB ID’s and oligomeric state of all 34 proteins in our validation dataset
3. benchmark_pdbs.csv: a csv file containing the PDB ID’s and oligomeric state of all 227 proteins in our training dataset
4. experimental_pdbs.csv: a csv file containing the PDB ID’s and oligomeric state of all 20 proteins in our training dataset
5. example_config.py: an example configuration file for inference with the SIDFold network.
6. model_5_multimer_v3.npz: the model weights file for the SIDFold network
