## Supplementary material for "Incorporating Surfaced-Induced Dissociation Mass Spectrometry Data into an AlphaFold-derived deep learning network improves protein structure prediction": example_config.py

```
import re
import copy
import importlib
import ml_collections as mlc


def set_inf(c, inf):
    for k, v in c.items():
        if isinstance(v, mlc.ConfigDict):
            set_inf(v, inf)
        elif k == "inf":
            c[k] = inf


def enforce_config_constraints(config):
    def string_to_setting(s):
        path = s.split('.')
        setting = config
        for p in path:
            setting = setting.get(p)

        return setting

    mutually_exclusive_bools = [
        (
            "model.template.average_templates", 
            "model.template.offload_templates"
        ),
        (
            "globals.use_lma",
            "globals.use_flash",
            "globals.use_deepspeed_evo_attention"
        ),
    ]

    for options in mutually_exclusive_bools:
        option_settings = [string_to_setting(o) for o in options]
        if sum(option_settings) > 1:
            raise ValueError(f"Only one of {', '.join(options)} may be set at a time")

    fa_is_installed = importlib.util.find_spec("flash_attn") is not None
    if config.globals.use_flash and not fa_is_installed:
        raise ValueError("use_flash requires that FlashAttention is installed")

    deepspeed_is_installed = importlib.util.find_spec("deepspeed") is not None
    ds4s_is_installed = deepspeed_is_installed and importlib.util.find_spec(
        "deepspeed.ops.deepspeed4science") is not None
    if config.globals.use_deepspeed_evo_attention and not ds4s_is_installed:
        raise ValueError(
            "use_deepspeed_evo_attention requires that DeepSpeed be installed "
            "and that the deepspeed.ops.deepspeed4science package exists"
        )

    if(
        config.globals.offload_inference and 
        not config.model.template.average_templates
    ):
        config.model.template.offload_templates = True


def model_config(
    name, 
    train=False, 
    low_prec=False, 
    long_sequence_inference=False,
    use_deepspeed_evoformer_attention=False,
):
    c = copy.deepcopy(config)
    # TRAINING PRESETS
    if name == "initial_training":
        # AF2 Suppl. Table 4, "initial training" setting
        pass
    elif name == "finetuning":
        # AF2 Suppl. Table 4, "finetuning" setting
        c.data.train.crop_size = 384
        c.data.train.max_extra_msa = 5120
        c.data.train.max_msa_clusters = 512
        c.loss.violation.weight = 1.
        c.loss.experimentally_resolved.weight = 0.01
    elif name == "finetuning_ptm":
        c.data.train.max_extra_msa = 5120
        c.data.train.crop_size = 384
        c.data.train.max_msa_clusters = 512
        c.loss.violation.weight = 1.
        c.loss.experimentally_resolved.weight = 0.01
        c.model.heads.tm.enabled = True
        c.loss.tm.weight = 0.1
    elif name == "finetuning_no_templ":
        # AF2 Suppl. Table 4, "finetuning" setting
        c.data.train.crop_size = 384
        c.data.train.max_extra_msa = 5120
        c.data.train.max_msa_clusters = 512
        c.model.template.enabled = False
        c.loss.violation.weight = 1.
        c.loss.experimentally_resolved.weight = 0.01
    elif name == "finetuning_no_templ_ptm":
        # AF2 Suppl. Table 4, "finetuning" setting
        c.data.train.crop_size = 384
        c.data.train.max_extra_msa = 5120
        c.data.train.max_msa_clusters = 512
        c.model.template.enabled = False
        c.loss.violation.weight = 1.
        c.loss.experimentally_resolved.weight = 0.01
        c.model.heads.tm.enabled = True
        c.loss.tm.weight = 0.1
    # INFERENCE PRESETS
    elif name == "model_1":
        # AF2 Suppl. Table 5, Model 1.1.1
        c.data.train.max_extra_msa = 5120
        c.data.predict.max_extra_msa = 5120
        c.data.common.reduce_max_clusters_by_max_templates = True
        c.data.common.use_templates = True
        c.data.common.use_template_torsion_angles = True
        c.model.template.enabled = True
    elif name == "model_2":
        # AF2 Suppl. Table 5, Model 1.1.2
        c.data.common.reduce_max_clusters_by_max_templates = True
        c.data.common.use_templates = True
        c.data.common.use_template_torsion_angles = True
        c.model.template.enabled = True
    elif name == "model_3":
        # AF2 Suppl. Table 5, Model 1.2.1
        c.data.train.max_extra_msa = 5120
        c.data.predict.max_extra_msa = 5120
        c.model.template.enabled = False
    elif name == "model_4":
        # AF2 Suppl. Table 5, Model 1.2.2
        c.data.train.max_extra_msa = 5120
        c.data.predict.max_extra_msa = 5120
        c.model.template.enabled = False
    elif name == "model_5":
        # AF2 Suppl. Table 5, Model 1.2.3
        c.model.template.enabled = False
    elif name == "model_1_ptm":
        c.data.train.max_extra_msa = 5120
        c.data.predict.max_extra_msa = 5120 
        c.data.common.reduce_max_clusters_by_max_templates = True
        c.data.common.use_templates = True
        c.data.common.use_template_torsion_angles = True
        c.model.template.enabled = True
        c.model.heads.tm.enabled = True
        c.loss.tm.weight = 0.1
    elif name == "model_2_ptm":
        c.data.common.reduce_max_clusters_by_max_templates = True
        c.data.common.use_templates = True
        c.data.common.use_template_torsion_angles = True
        c.model.template.enabled = True
        c.model.heads.tm.enabled = True
        c.loss.tm.weight = 0.1
    elif name == "model_3_ptm":
        c.data.train.max_extra_msa = 5120
        c.data.predict.max_extra_msa = 5120
        c.model.template.enabled = False
        c.model.heads.tm.enabled = True
        c.loss.tm.weight = 0.1
    elif name == "model_4_ptm":
        c.data.train.max_extra_msa = 5120
        c.data.predict.max_extra_msa = 5120
        c.model.template.enabled = False
        c.model.heads.tm.enabled = True
        c.loss.tm.weight = 0.1
    elif name == "model_5_ptm":
        c.model.template.enabled = False
        c.model.heads.tm.enabled = True
        c.loss.tm.weight = 0.1
    elif name.startswith("seq"):  # SINGLE SEQUENCE EMBEDDING PRESETS
        c.update(seq_mode_config.copy_and_resolve_references())
        if name == "seqemb_initial_training":
            c.data.train.max_msa_clusters = 1
            c.data.eval.max_msa_clusters = 1
            c.data.train.block_delete_msa = False
            c.data.train.max_distillation_msa_clusters = 1
        elif name == "seqemb_finetuning":
            c.data.train.max_msa_clusters = 1
            c.data.eval.max_msa_clusters = 1
            c.data.train.block_delete_msa = False
            c.data.train.max_distillation_msa_clusters = 1
            c.data.train.crop_size = 384
            c.loss.violation.weight = 1.
            c.loss.experimentally_resolved.weight = 0.01
        elif name == "seq_model_esm1b":
            c.data.common.use_templates = True
            c.data.common.use_template_torsion_angles = True
            c.model.template.enabled = True
            c.data.predict.max_msa_clusters = 1
        elif name == "seq_model_esm1b_ptm":
            c.data.common.use_templates = True
            c.data.common.use_template_torsion_angles = True
            c.model.template.enabled = True
            c.data.predict.max_msa_clusters = 1
            c.model.heads.tm.enabled = True
            c.loss.tm.weight = 0.1
    elif "multimer" in name:  # MULTIMER PRESETS
        c.update(multimer_config_update.copy_and_resolve_references())

        # Not used in multimer
        del c.model.template.template_pointwise_attention
        del c.loss.fape.backbone

        # TODO: Change max_msa_clusters and max_extra_msa to multimer feats within model
        if re.fullmatch("^model_[1-5]_multimer(_v2)?$", name):
            #c.model.input_embedder.num_msa = 252
            #c.model.extra_msa.extra_msa_embedder.num_extra_msa = 1152
            c.data.train.crop_size = 384

            c.data.train.max_msa_clusters = 252
            c.data.eval.max_msa_clusters = 252
            c.data.predict.max_msa_clusters = 252

            c.data.train.max_extra_msa = 1152
            c.data.eval.max_extra_msa = 1152
            c.data.predict.max_extra_msa = 1152

            c.model.evoformer_stack.fuse_projection_weights = False
            c.model.extra_msa.extra_msa_stack.fuse_projection_weights = False
            c.model.template.template_pair_stack.fuse_projection_weights = False
        elif name == 'model_4_multimer_v3':
            #c.model.extra_msa.extra_msa_embedder.num_extra_msa = 1152
            c.data.train.max_extra_msa = 1152
            c.data.eval.max_extra_msa = 1152
            c.data.predict.max_extra_msa = 1152
        elif name == 'model_5_multimer_v3':
            #c.model.extra_msa.extra_msa_embedder.num_extra_msa = 1152
            c.data.train.max_extra_msa = 1152 #RMB changed, default is 1152
            c.data.eval.max_extra_msa = 1152 #RMB changed, default is 1152
            c.data.train.max_extra_msa = 5120 #RMB added from finetuning
            c.data.predict.max_extra_msa = 1152 #RMB changed, default is 1152
            c.data.train.crop_size = 384 #RMB added this line, default is 384
            #c.data.eval.crop_size = 1400 #RMB added this line, default is 384, DO NOT TURN THIS ON, IT WILL MESS UP THE LOSS
            #c.model.template.enabled = False #RMB added this line in from model_5 monomer config
            #c.data.common.use_templates = False #RMB added from inference config 1
            #c.loss.violation.weight = 1. #RMB added from finetuning
            c.loss.tm.weight = 0.1 #RMB added from ptm
            #c.model.template.enabled = False #RMB added from ptm
            #c.model.heads.tm.enabled = True #RMB added from ptm
    else:
        raise ValueError("Invalid model name")

    if long_sequence_inference:
        assert(not train) 
        c.globals.offload_inference = True
        # Default to DeepSpeed memory-efficient attention kernel unless use_lma is explicitly set
        c.globals.use_deepspeed_evo_attention = True if not c.globals.use_lma else False
        c.globals.use_flash = False
        c.model.template.offload_inference = True
        c.model.template.template_pair_stack.tune_chunk_size = False
        c.model.extra_msa.extra_msa_stack.tune_chunk_size = False
        c.model.evoformer_stack.tune_chunk_size = False
    
    if use_deepspeed_evoformer_attention:
        c.globals.use_deepspeed_evo_attention = True 
    
    if train:
        c.globals.blocks_per_ckpt = 1
        c.globals.chunk_size = None
        c.globals.use_lma = False
        c.globals.offload_inference = False #RMB changed to True, default is False
        c.model.template.average_templates = False
        c.model.template.offload_templates = False #RMB changed to True, default is False
    
    if low_prec:
        c.globals.eps = 1e-4
        # If we want exact numerical parity with the original, inf can't be
        # a global constant
        set_inf(c, 1e4)

    enforce_config_constraints(c)

    return c


c_z = mlc.FieldReference(128, field_type=int)
c_m = mlc.FieldReference(256, field_type=int)
c_t = mlc.FieldReference(64, field_type=int)
c_e = mlc.FieldReference(64, field_type=int)
c_s = mlc.FieldReference(384, field_type=int)
c_b = mlc.FieldReference(8, field_type=int) #RMB added from ED config

# For seqemb mode, dimension size of the per-residue sequence embedding passed to the model
# In current model, the dimension size is the ESM-1b dimension size i.e. 1280.
preemb_dim_size = mlc.FieldReference(1280, field_type=int)

blocks_per_ckpt = mlc.FieldReference(None, field_type=int)
chunk_size = mlc.FieldReference(4, field_type=int)
aux_distogram_bins = mlc.FieldReference(64, field_type=int)
tm_enabled = mlc.FieldReference(False, field_type=bool)
eps = mlc.FieldReference(1e-8, field_type=float)
templates_enabled = mlc.FieldReference(True, field_type=bool) 
embed_template_torsion_angles = mlc.FieldReference(True, field_type=bool)
tune_chunk_size = mlc.FieldReference(True, field_type=bool)

NUM_RES = "num residues placeholder"
NUM_MSA_SEQ = "msa placeholder"
NUM_EXTRA_SEQ = "extra msa placeholder"
NUM_TEMPLATES = "num templates placeholder"

config = mlc.ConfigDict(
    {
        "data": {
            "common": {
                "feat": {
                    "aatype": [NUM_RES],
                    "all_atom_mask": [NUM_RES, None],
                    "all_atom_positions": [NUM_RES, None, None],
                    "alt_chi_angles": [NUM_RES, None],
                    "atom14_alt_gt_exists": [NUM_RES, None],
                    "atom14_alt_gt_positions": [NUM_RES, None, None],
                    "atom14_atom_exists": [NUM_RES, None],
                    "atom14_atom_is_ambiguous": [NUM_RES, None],
                    "atom14_gt_exists": [NUM_RES, None],
                    "atom14_gt_positions": [NUM_RES, None, None],
                    "atom37_atom_exists": [NUM_RES, None],
                    "backbone_rigid_mask": [NUM_RES],
                    "backbone_rigid_tensor": [NUM_RES, None, None],
                    "bert_mask": [NUM_MSA_SEQ, NUM_RES],
                    "chi_angles_sin_cos": [NUM_RES, None, None],
                    "chi_mask": [NUM_RES, None],
                    "extra_deletion_value": [NUM_EXTRA_SEQ, NUM_RES],
                    "extra_has_deletion": [NUM_EXTRA_SEQ, NUM_RES],
                    "extra_msa": [NUM_EXTRA_SEQ, NUM_RES],
                    "extra_msa_mask": [NUM_EXTRA_SEQ, NUM_RES],
                    "extra_msa_row_mask": [NUM_EXTRA_SEQ],
                    "is_distillation": [],
                    "msa_feat": [NUM_MSA_SEQ, NUM_RES, None],
                    "msa_mask": [NUM_MSA_SEQ, NUM_RES],
                    "msa_row_mask": [NUM_MSA_SEQ],
                    "no_recycling_iters": [],
                    "pseudo_beta": [NUM_RES, None],
                    "pseudo_beta_mask": [NUM_RES],
                    "residue_index": [NUM_RES],
                    "residx_atom14_to_atom37": [NUM_RES, None],
                    "residx_atom37_to_atom14": [NUM_RES, None],
                    "resolution": [],
                    "rigidgroups_alt_gt_frames": [NUM_RES, None, None, None],
                    "rigidgroups_group_exists": [NUM_RES, None],
                    "rigidgroups_group_is_ambiguous": [NUM_RES, None],
                    "rigidgroups_gt_exists": [NUM_RES, None],
                    "rigidgroups_gt_frames": [NUM_RES, None, None, None],
                    "seq_length": [],
                    "seq_mask": [NUM_RES],
                    "target_feat": [NUM_RES, None],
                    "template_aatype": [NUM_TEMPLATES, NUM_RES],
                    "template_all_atom_mask": [NUM_TEMPLATES, NUM_RES, None],
                    "template_all_atom_positions": [
                        NUM_TEMPLATES, NUM_RES, None, None,
                    ],
                    "template_alt_torsion_angles_sin_cos": [
                        NUM_TEMPLATES, NUM_RES, None, None,
                    ],
                    "template_backbone_rigid_mask": [NUM_TEMPLATES, NUM_RES],
                    "template_backbone_rigid_tensor": [
                        NUM_TEMPLATES, NUM_RES, None, None,
                    ],
                    "template_mask": [NUM_TEMPLATES],
                    "template_pseudo_beta": [NUM_TEMPLATES, NUM_RES, None],
                    "template_pseudo_beta_mask": [NUM_TEMPLATES, NUM_RES],
                    "template_sum_probs": [NUM_TEMPLATES, None],
                    "template_torsion_angles_mask": [
                        NUM_TEMPLATES, NUM_RES, None,
                    ],
                    "template_torsion_angles_sin_cos": [
                        NUM_TEMPLATES, NUM_RES, None, None,
                    ],
                    "true_msa": [NUM_MSA_SEQ, NUM_RES],
                    "use_clamped_fape": [],
                },
                "block_delete_msa": {
                    "msa_fraction_per_block": 0.3,
                    "randomize_num_blocks": False,
                    "num_blocks": 5,
                },
                "masked_msa": {
                    "profile_prob": 0.1,
                    "same_prob": 0.1,
                    "uniform_prob": 0.1,
                },
                "max_recycling_iters": 3,
                "msa_cluster_features": True,
                "reduce_msa_clusters_by_max_templates": False,
                "resample_msa_in_recycling": True,
                "template_features": [
                    "template_all_atom_positions",
                    "template_sum_probs",
                    "template_aatype",
                    "template_all_atom_mask",
                ],
                "unsupervised_features": [
                    "aatype",
                    "residue_index",
                    "msa",
                    "num_alignments",
                    "seq_length",
                    "between_segment_residues",
                    "deletion_matrix",
                    "no_recycling_iters",
                ],
                "use_templates": templates_enabled,
                "use_template_torsion_angles": embed_template_torsion_angles,
            },
            "seqemb_mode": { # Configuration for sequence embedding mode
                "enabled": False, # If True, use seq emb instead of MSA
            },
            "supervised": {
                "clamp_prob": 0.9,
                "supervised_features": [
                    "all_atom_mask",
                    "all_atom_positions",
                    "resolution",
                    "use_clamped_fape",
                    "is_distillation",
                ],
            },
            "predict": {
                "fixed_size": True,
                "subsample_templates": False,  # We want top templates.
                "block_delete_msa": False,
                "masked_msa_replace_fraction": 0.15,
                "max_msa_clusters": 512,
                "max_extra_msa": 1024,
                "max_template_hits": 4,
                "max_templates": 4,
                "crop": False,
                "crop_size": None,
                "spatial_crop_prob": None,
                "interface_threshold": None,
                "supervised": False,
                "uniform_recycling": False,
            },
            "eval": {
                "fixed_size": True,
                "subsample_templates": False,  #RMB changed from false
                "block_delete_msa": False, #RMB changed from false
                "masked_msa_replace_fraction": 0.15,
                "max_msa_clusters": 128, #RMB changed, default is 128
                "max_extra_msa": 1024, #RMB changed, default is 1024
                "max_template_hits": 4,
                "max_templates": 4,
                "crop": False,
                "crop_size": None,
                "spatial_crop_prob": None,
                "interface_threshold": None,
                "supervised": True,
                "uniform_recycling": False,
            },
            "train": {
                "fixed_size": True,
                "subsample_templates": False, #RMB changed from True
                "block_delete_msa": True,
                "masked_msa_replace_fraction": 0.15,
                "max_msa_clusters": 128, #RMB changed, default is 128
                "max_extra_msa": 1024, #RMB changed, default is 1024
                "max_template_hits": 4,
                "max_templates": 4,
                "shuffle_top_k_prefiltered": 20,
                "crop": True,
                "crop_size": 256, #RMB changed to 384
                "spatial_crop_prob": 0.,
                "interface_threshold": None,
                "supervised": True,
                "clamp_prob": 0.9,
                "max_distillation_msa_clusters": 1000,
                "uniform_recycling": True,
                "distillation_prob": 0.75,
            },
            "data_module": {
                "use_small_bfd": True,
                "data_loaders": {
                    "batch_size": 1,
                    "num_workers": 20, #RMB set this to 20 for real training
                    "pin_memory": False, #RMB changed to False
                },
            },
        },
        # Recurring FieldReferences that can be changed globally here
        "globals": {
            "blocks_per_ckpt": blocks_per_ckpt,
            "chunk_size": chunk_size,
            # Use DeepSpeed memory-efficient attention kernel. Mutually
            # exclusive with use_lma and use_flash.
            "use_deepspeed_evo_attention": False, #RMB default is False, tried setting to True (also tried use_flash)
            # Use Staats & Rabe's low-memory attention algorithm. Mutually
            # exclusive with use_deepspeed_evo_attention and use_flash.
            "use_lma": False,
            # Use FlashAttention in selected modules. Mutually exclusive with 
            # use_deepspeed_evo_attention and use_lma. Doesn't work that well
            # on long sequences (>1000 residues).
            "use_flash": False, #RMB default is False, tried setting to True (also tried deepspeed_evo_attention)
            "offload_inference": False,
            "c_z": c_z,
            "c_m": c_m,
            "c_t": c_t,
            "c_e": c_e,
            "c_s": c_s,
            "eps": eps,
            "is_multimer": False,
            "seqemb_mode_enabled": False, # Global flag for enabling seq emb mode
        },
        "model": {
            "_mask_trans": False,
            "input_embedder": {
                "tf_dim": 22,
                "msa_dim": 49,
                "c_z": c_z,
                "c_m": c_m,
                "relpos_k": 32,
            },
            "recycling_embedder": {
                "c_z": c_z,
                "c_m": c_m,
                "min_bin": 3.25,
                "max_bin": 20.75,
                "no_bins": 15,
                "inf": 1e8,
            },
            "template": {
                "distogram": {
                    "min_bin": 3.25,
                    "max_bin": 50.75,
                    "no_bins": 39,
                },
                "template_single_embedder": {
                    # DISCREPANCY: c_in is supposed to be 51.
                    "c_in": 57,
                    "c_out": c_m,
                },
                "template_pair_embedder": {
                    "c_in": 88,
                    "c_out": c_t,
                },
                "template_pair_stack": {
                    "c_t": c_t,
                    # DISCREPANCY: c_hidden_tri_att here is given in the supplement
                    # as 64. In the code, it's 16.
                    "c_hidden_tri_att": 16,
                    "c_hidden_tri_mul": 64,
                    "no_blocks": 2,
                    "no_heads": 4,
                    "pair_transition_n": 2,
                    "dropout_rate": 0.25,
                    "tri_mul_first": False,
                    "fuse_projection_weights": False,
                    "blocks_per_ckpt": blocks_per_ckpt,
                    "tune_chunk_size": tune_chunk_size,
                    "inf": 1e9,
                },
                "template_pointwise_attention": {
                    "c_t": c_t,
                    "c_z": c_z,
                    # DISCREPANCY: c_hidden here is given in the supplement as 64.
                    # It's actually 16.
                    "c_hidden": 16,
                    "no_heads": 4,
                    "inf": 1e5,  # 1e9,
                },
                "inf": 1e5,  # 1e9,
                "eps": eps,  # 1e-6,
                "enabled": templates_enabled,
                "embed_angles": embed_template_torsion_angles,
                "use_unit_vector": False,
                # Approximate template computation, saving memory.
                # In our experiments, results are equivalent to or better than
                # the stock implementation. Should be enabled for all new
                # training runs.
                "average_templates": False,
                # Offload template embeddings to CPU memory. Vastly reduced
                # memory consumption at the cost of a modest increase in
                # runtime. Useful for inference on very long sequences.
                # Mutually exclusive with average_templates. Automatically
                # enabled if offload_inference is set.
                "offload_templates": False,
            },
            "extra_msa": {
                "extra_msa_embedder": {
                    "c_in": 25,
                    "c_out": c_e,
                },
                "extra_msa_stack": {
                    "c_m": c_e,
                    "c_z": c_z,
                    "c_hidden_msa_att": 8,
                    "c_hidden_opm": 32,
                    "c_hidden_mul": 128,
                    "c_hidden_pair_att": 32,
                    "no_heads_msa": 8,
                    "no_heads_pair": 4,
                    "no_blocks": 4,
                    "transition_n": 4,
                    "msa_dropout": 0.15,
                    "pair_dropout": 0.25,
                    "opm_first": False,
                    "fuse_projection_weights": False,
                    "clear_cache_between_blocks": False,
                    "tune_chunk_size": tune_chunk_size,
                    "inf": 1e9,
                    "eps": eps,  # 1e-10,
                    "ckpt": blocks_per_ckpt is not None,
                },
                "enabled": True,
            },
            "evoformer_stack": {
                "c_m": c_m,
                "c_z": c_z,
                "c_hidden_msa_att": 32,
                "c_hidden_opm": 32,
                "c_hidden_mul": 128,
                "c_hidden_pair_att": 32,
                "c_s": c_s,
                "no_heads_msa": 8,
                "no_heads_pair": 4,
                "no_blocks": 48,
                "transition_n": 4,
                "msa_dropout": 0.15,
                "pair_dropout": 0.25,
                "no_column_attention": False,
                "opm_first": False,
                "fuse_projection_weights": False,
                "blocks_per_ckpt": blocks_per_ckpt,
                "clear_cache_between_blocks": False,
                "tune_chunk_size": tune_chunk_size,
                "inf": 1e9,
                "eps": eps,  # 1e-10,
            },
            "structure_module": {
                "c_s": c_s,
                "c_z": c_z,
                "c_ipa": 16,
                "c_resnet": 128,
                "no_heads_ipa": 12,
                "no_qk_points": 4,
                "no_v_points": 8,
                "dropout_rate": 0.1,
                "no_blocks": 8,
                "no_transition_layers": 1,
                "no_resnet_blocks": 2,
                "no_angles": 7,
                "trans_scale_factor": 10,
                "epsilon": eps,  # 1e-12,
                "inf": 1e5,
            },
            "heads": {
                "lddt": {
                    "no_bins": 50,
                    "c_in": c_s,
                    "c_hidden": 128,
                },
                "distogram": {
                    "c_z": c_z,
                    "no_bins": aux_distogram_bins,
                },
                "tm": {
                    "c_z": c_z,
                    "no_bins": aux_distogram_bins,
                    "enabled": tm_enabled,
                },
                "masked_msa": {
                    "c_m": c_m,
                    "c_out": 23,
                },
                "experimentally_resolved": {
                    "c_s": c_s,
                    "c_out": 37,
                },
            },
            # A negative value indicates that no early stopping will occur, i.e.
            # the model will always run `max_recycling_iters` number of recycling
            # iterations. A positive value will enable early stopping if the
            # difference in pairwise distances is less than the tolerance between
            # recycling steps.
            "recycle_early_stop_tolerance": -1.
        },
        "relax": {
            "max_iterations": 0,  # no max
            "tolerance": 10.0,
            "stiffness": 10.0,
            "max_outer_iterations": 20,
            "exclude_residues": [],
        },
        "loss": {
            "distogram": {
                "min_bin": 2.3125,
                "max_bin": 21.6875,
                "no_bins": 64,
                "eps": eps,  # 1e-6,
                "weight": 0.3,
            },
            "experimentally_resolved": {
                "eps": eps,  # 1e-8,
                "min_resolution": 0.1,
                "max_resolution": 3.0,
                "weight": 0.0,
            },
            "fape": {
                "backbone": {
                    "clamp_distance": 10.0,
                    "loss_unit_distance": 10.0,
                    "weight": 0.5,
                },
                "sidechain": {
                    "clamp_distance": 10.0,
                    "length_scale": 10.0,
                    "weight": 0.5,
                },
                "eps": 1e-4,
                "weight": 1.0,
            },
            "plddt_loss": {
                "min_resolution": 0.1,
                "max_resolution": 3.0,
                "cutoff": 15.0,
                "no_bins": 50,
                "eps": eps,  # 1e-10,
                "weight": 0.01,
            },
            "masked_msa": {
                "num_classes": 23,
                "eps": eps,  # 1e-8,
                "weight": 2.0,
            },
            "supervised_chi": {
                "chi_weight": 0.5,
                "angle_norm_weight": 0.01,
                "eps": eps,  # 1e-6,
                "weight": 1.0,
            },
            "violation": {
                "violation_tolerance_factor": 12.0,
                "clash_overlap_tolerance": 1.5,
                "average_clashes": False,
                "eps": eps,  # 1e-6,
                "weight": 0.0,
            },
            "tm": {
                "max_bin": 31,
                "no_bins": 64,
                "min_resolution": 0.1,
                "max_resolution": 3.0,
                "eps": eps,  # 1e-8,
                "weight": 0.,
                "enabled": tm_enabled,
            },
            "chain_center_of_mass": {
                "clamp_distance": -4.0,
                "weight": 0.,
                "eps": eps,
                "enabled": False,
            },
            "eps": eps,
        },
        "ema": {"decay": 0.999},
    }
)

multimer_config_update = mlc.ConfigDict({
    "globals": {
        #"use_deepspeed_evo_attention": False, #RMB added this line 
        #"offload_inference": True, #RMB added this line
        "is_multimer": True
    },
    "data": {
        "common": {
            "feat": {
                "aatype": [NUM_RES],
                "all_atom_mask": [NUM_RES, None],
                "all_atom_positions": [NUM_RES, None, None],
                # "all_chains_entity_ids": [],  # TODO: Resolve missing features, remove processed msa feats
                # "all_crops_all_chains_mask": [],
                # "all_crops_all_chains_positions": [],
                # "all_crops_all_chains_residue_ids": [],
                "assembly_num_chains": [],
                "asym_id": [NUM_RES],
                "atom14_atom_exists": [NUM_RES, None],
                "atom37_atom_exists": [NUM_RES, None],
                "bert_mask": [NUM_MSA_SEQ, NUM_RES],
                "cluster_bias_mask": [NUM_MSA_SEQ],
                "cluster_profile": [NUM_MSA_SEQ, NUM_RES, None],
                "cluster_deletion_mean": [NUM_MSA_SEQ, NUM_RES],
                "deletion_matrix": [NUM_MSA_SEQ, NUM_RES],
                "deletion_mean": [NUM_RES],
                "entity_id": [NUM_RES],
                "entity_mask": [NUM_RES],
                "extra_deletion_matrix": [NUM_EXTRA_SEQ, NUM_RES],
                "extra_msa": [NUM_EXTRA_SEQ, NUM_RES],
                "extra_msa_mask": [NUM_EXTRA_SEQ, NUM_RES],
                # "mem_peak": [],
                "msa": [NUM_MSA_SEQ, NUM_RES],
                "msa_feat": [NUM_MSA_SEQ, NUM_RES, None],
                "msa_mask": [NUM_MSA_SEQ, NUM_RES],
                "msa_profile": [NUM_RES, None],
                "num_alignments": [],
                "num_templates": [],
                # "queue_size": [],
                "residue_index": [NUM_RES],
                "residx_atom14_to_atom37": [NUM_RES, None],
                "residx_atom37_to_atom14": [NUM_RES, None],
                "resolution": [],
                "seq_length": [],
                "seq_mask": [NUM_RES],
                "sym_id": [NUM_RES],
                "target_feat": [NUM_RES, None],
                "template_aatype": [NUM_TEMPLATES, NUM_RES],
                "template_all_atom_mask": [NUM_TEMPLATES, NUM_RES, None],
                "template_all_atom_positions": [
                    NUM_TEMPLATES, NUM_RES, None, None,
                ],
                "true_msa": [NUM_MSA_SEQ, NUM_RES],
                "erms_data": [None], #RMB added
                "erms_sim": [None], #RMB added
                "name": [None] #RMB added
            },
            "max_recycling_iters": 3,  # For training, value is 3, RMB changed to 3 for training
            "unsupervised_features": [
                    "aatype",
                    "residue_index",
                    "msa",
                    "num_alignments",
                    "seq_length",
                    "between_segment_residues",
                    "deletion_matrix",
                    "no_recycling_iters",
                    # Additional multimer features
                    "msa_mask",
                    "seq_mask",
                    "asym_id",
                    "entity_id",
                    "sym_id",
                    "erms_data", #RMB added
                    "erms_sim", #RMB added
                    "name", #RMB added
                ]
        },
        "supervised": {
            "clamp_prob": 1.
        },
        # TODO: Change max_msa_clusters and max_extra_msa to multimer feats within model:
        # c.model.input_embedder.num_msa = 508
        # c.model.extra_msa.extra_msa_embedder.num_extra_msa = 2048
        "predict": {
            "max_msa_clusters": 508,
            "max_extra_msa": 0
        },
        "eval": {
            "max_msa_clusters": 508, #RMB changed, default value is 508
            "max_extra_msa": 2048 #RMB changed, default value is 2048
        },
        "train": {
            "max_msa_clusters": 508, #RMB changed, default value is 508
            "max_extra_msa": 2048, #RMB changed, default value is 2048
            "block_delete_msa" : False, #RMB changed to True, default is False
            "crop_size": 640, #RMB changed, default value is 640
            "spatial_crop_prob": 0.5,
            "interface_threshold": 10.,
            "clamp_prob": 1.,
        },
    },
    "model": {
        "input_embedder": {
            "tf_dim": 21,
            #"num_msa": 508,
            "max_relative_chain": 2,
            "max_relative_idx": 32,
            "use_chain_relative": True
        },
        "template": {
            "template_single_embedder": {
                "c_in": 34,
                "c_out": c_m
            },
            "template_pair_embedder": {
                "c_in": c_z,
                "c_out": c_t,
                "c_dgram": 39,
                "c_aatype": 22
            },
            "template_pair_stack": {
                "tri_mul_first": True,
                "fuse_projection_weights": True
            },
            "c_t": c_t,
            "c_z": c_z,
            "use_unit_vector": True
        },
        "extra_msa": {
            # "extra_msa_embedder": {
            #     "num_extra_msa": 2048
            # },
            "extra_msa_stack": { #RMB commented out this extra_msa_stack include
                "opm_first": True,
                "fuse_projection_weights": True
            }
        },
        "evoformer_stack": {
            "opm_first": True,
            "fuse_projection_weights": True
        },
        "structure_module": {
            "trans_scale_factor": 20
        },
        "heads": {
            "tm": {
                "ptm_weight": 0.2,
                "iptm_weight": 0.8,
                "enabled": True
            },
            "masked_msa": {
                "c_out": 22
            },
        },
        "recycle_early_stop_tolerance": -1,  # For training, value is -1. #RMB changed to -1 for training, default is 0.5
        "relax": { #RMB needs to add relax here to use in model.py, must be in model section of multimer config to be called correctly
                "max_iterations": 0,  # no max
                "tolerance": 10.0,
                "stiffness": 10.0,
                "max_outer_iterations": 20,
                "exclude_residues": [],
        },
    },
    "loss": {
        "fape": {
            "intra_chain_backbone": {
                "clamp_distance": 10.0,
                "loss_unit_distance": 10.0,
                "weight": 0.5
            },
            "interface_backbone": {
                "clamp_distance": 30.0,
                "loss_unit_distance": 20.0,
                "weight": 0.5
            }
        },
        "masked_msa": {
            "num_classes": 22
        },
        "erms": { #RMB added for ERMS loss function, current weight is arbitrary
            "weight": 0.01,  
        },
        "violation": {
            "average_clashes": True,
            "weight": 0.03 # Not finetuning
        },
        "tm": {
            "weight": 0.1,
            "enabled": True
        },
        "chain_center_of_mass": {
            "weight": 0.05,
            "enabled": True
        }
    }
})


seq_mode_config = mlc.ConfigDict({
    "data": {
        "common": {
            "feat": {
                "seq_embedding": [NUM_RES, None],
            },
            "seqemb_features": [ # List of features to be generated in seqemb mode
                "seq_embedding"
            ],
        },
        "seqemb_mode": { # Configuration for sequence embedding mode
            "enabled": True, # If True, use seq emb instead of MSA
        },
    },
    "globals": {
        "seqemb_mode_enabled": True,
    },
    "model": {
        "preembedding_embedder": {  # Used in sequence embedding mode
            "tf_dim": 22,
            "preembedding_dim": preemb_dim_size,
            "c_z": c_z,
            "c_m": c_m,
            "relpos_k": 32,
        },
        "extra_msa": {
            "enabled": False  # Disable Extra MSA Stack
        },
        "evoformer_stack": {
            "no_column_attention": True  # Turn off Evoformer's column attention
        },
    }
})
```
